# TREND: A generalizable synthetic enhancer discovery platform for targeted immunotherapy

**DOI:** 10.64898/2025.12.08.693012

**Authors:** Yingbo Jia, Chu-Yen Chen, Bo Zhu, Zhigui Wu, Yi-Chia Wu, Renqi Wang, Abdulmajeed I. Salamah, Sushmita Halder, Yen-Nien Liu, Yingjie Guo, Ying Chen, Luca Scimeca, Hang Yin, Tejas Sabu, Yutao Wang, Enoch B. Antwi, Yang Wang, Ming-Ru Wu

## Abstract

Synthetic enhancers with high specificity are crucial for therapeutic gene control. However, experimental screens and machine learning-guided design typically require context-specific datasets, limiting generalizability. Because transcription factor (TF) activity reflects cellular state, TF-responsive enhancer libraries offer a universal starting point. Here, we developed TREND (transcription factor-responsive enhancer discovery), a massively parallel reporter assay of ∼2.7 million enhancer-barcode constructs representing 57,715 designs. TREND covers 1,068 motif-annotated proteins, including 729 confirmed TFs across 49 DNA-binding domain families. Applied to ovarian cancer, TREND identified enhancers that discriminate cancer from normal epithelial cells. These enhancers enabled protein-interaction-based AND-gate circuits with reduced OFF-state leakage and amplified ON-state output, driving tumor-restricted expression of combinatorial immune effectors and robust antitumor responses in murine ovarian cancer models. TREND also identified T-cell activation-responsive enhancers with greater inducibility and lower basal activity than conventional NFAT-motif-based elements. Together, TREND provides a generalizable framework for context-specific enhancer discovery and therapeutic gene regulation.

## Introduction

Enhancers orchestrate cell type- and state-specific transcriptional programs by integrating the activity of multiple transcription factors (TFs)^1,2^. As a result, enhancer activity reflects the underlying transcriptional state of a cell and can encode cell identity and regulatory programs^3,4^. This intrinsic logic makes enhancers powerful elements for engineering precise gene control. In most gene therapy applications, cell-type-specific enhancers are essential for achieving selective and safe transgene expression^5^. In cancer immunotherapy, restricting transgene expression to malignant cells maximizes therapeutic targeting while minimizing damage to normal tissues. Similarly, in engineered immune-cell therapies, confining gene expression to activated rather than resting immune cells ensures that effector functions are triggered only upon antigen engagement, preventing unintended systemic immune activation^6,7^.

Several high-throughput approaches have been developed to identify regulatory elements that drive context-specific gene expression. Methods such as massively parallel reporter assays (MPRAs) and self-transcribing active regulatory region sequencing (STARR-seq) enable large-scale functional profiling of candidate regulatory elements and have facilitated the discovery of enhancers active in specific cellular contexts^8–13^. However, these approaches generally interrogate genomic sequences, requiring candidate elements pre-identified in the cell type or system of interest. Their applicability is therefore constrained by available regulatory annotations and may not generalize across new cellular or disease contexts. In parallel, machine-learning models have been used to computationally design cell-type-specific synthetic enhancers, with successful validation in mammalian systems^14–16^. However, these strategies rely on high-quality training datasets from active regulatory regions in the target cell type, which are often limited to particular developmental or steady-state contexts. Applying them to new settings therefore requires generating additional datasets or retraining models. Collectively, both classes of approach depend on pre-existing regulatory annotations or context-specific training data rather than directly querying endogenous TF activity, limiting their generalizability across cellular contexts.

An alternative strategy is to directly leverage TF activity as a context-dependent regulatory signal. Because enhancer activity is ultimately determined by TF binding, synthetic enhancers composed of TF binding sites (TFBSs) can serve as reporters of endogenous TF activity. We previously demonstrated that synthetic enhancers constructed from tandem TFBS repeats can be screened to identify certain context-responsive regulatory elements^17^. However, that initial platform had three key limitations: (i) its motif collection covered only a fraction of known TF binding preferences; (ii) it relied on consensus sequences without affinity-modulating variants needed for threshold-dependent responses; and (iii) enhancer identification depended on iterative FACS-based enrichment, constraining scalability and precluding analysis of fragile or rapidly changing cell states.

Here, we developed TREND (transcription factor-responsive enhancer discovery), a scalable platform for systematic identification of TF-responsive synthetic enhancers. TREND comprises ∼2.7 million unique enhancer-barcode constructs converging to 57,715 enhancer designs, screened in parallel using a lentiviral MPRA-based framework. This enables direct and quantitative measurement of enhancer activity across diverse cellular contexts. The library covers 1,068 proteins with annotated DNA-binding motifs, of which 729 are confirmed TFs across 49 DNA-binding domain families. Using this platform, we identified cancer-selective and T-cell activation-responsive enhancers and demonstrated their utility in diverse synthetic biology applications, including logic-gated gene circuit design and in vivo cancer immunotherapy. These findings position TREND as a generalizable strategy for leveraging endogenous transcriptional programs to discover and deploy context-specific regulatory elements across biological systems.

## Results

### TREND platform identifies candidate cancer-selective synthetic enhancers

To enable high-throughput screening of synthetic enhancers, we developed TREND, a lentiviral-based MPRA framework that quantifies TF-responsive enhancer activity. To enhance transcriptional output and enable selective readout of individual TF activities, each synthetic enhancer was built from tandem repeats of a single TFBS. We assembled a library of position probability matrices (PPMs) and position weight matrices (PWMs) from 14 public databases, including JASPAR, CIS-BP, and HOCOMOCO, as well as from protein-DNA interaction screens^18^ (Figure 1A). From these motifs, we designed ∼2.7 million unique enhancer-barcode constructs, converging to 57,715 enhancer designs. These target 1,068 proteins with annotated DNA-binding motifs, of which 729 are confirmed sequence-specific transcription factors. Together, these TFs span 49 DNA-binding domain families including homeodomain, C2H2 zinc finger, bHLH, nuclear receptor, bZIP, and forkhead families^3^ (Figure 1B and 1C). The remaining 339 targets were derived from unbiased protein-DNA interaction screens and may mediate regulatory activity through non-canonical DNA-binding mechanisms^19^. For brevity, we refer to all 1,068 targets collectively as TFs hereafter. Notably, this library captures a substantial fraction of regulatory programs relevant to cancer and cell identity: 75% of candidate lineage-specific master transcription factors (MTFs) across 34 tumor types and 70% of core identity transcription factors across 233 cell and tissue types (Figure 1D, 1E)^3,20,21^. This breadth provides a versatile starting point for identifying context-specific regulatory elements across diverse cellular and disease settings.

**Figure 1.**
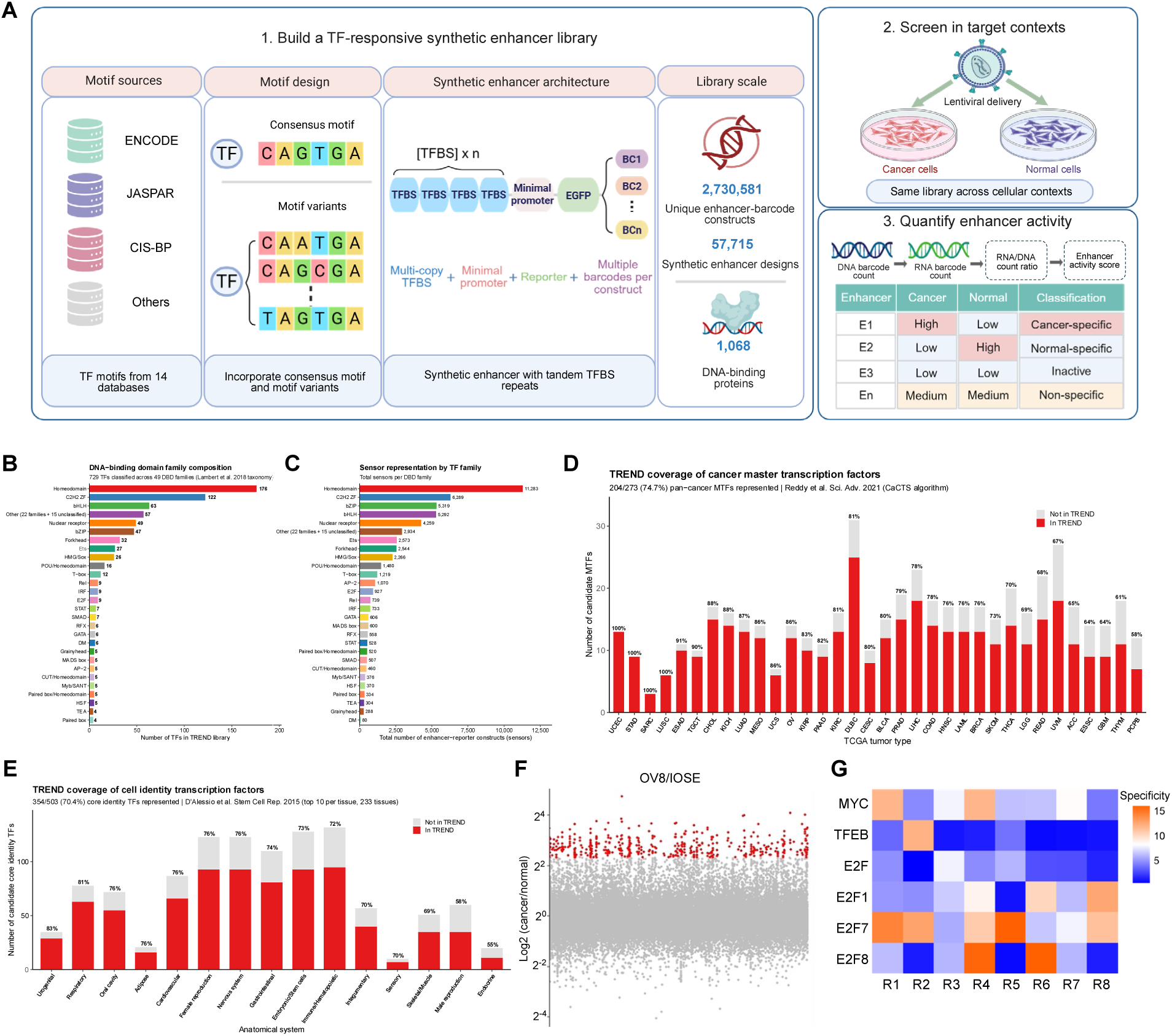
TREND platform for high-throughput discovery of synthetic enhancers. (A) Schematic of the TREND workflow. (1) PPMs and PWMs from 1,068 proteins with annotated DNA-binding motifs (14 databases) were used to generate 2,730,581 (∼2.7 million) unique enhancer-barcode constructs spanning 57,715 enhancer designs, each comprising consensus sequences and nine high-probability variants per matrix. (2) Each sequence was arrayed in tandem upstream of an adenoviral minimal promoter with a linked DNA barcode, packaged into a pooled lentiviral library, and delivered into target cells. (3) Enhancer activity was quantified by targeted NGS of barcode-linked RNA and DNA, with activity and specificity inferred from RNA/DNA ratios across conditions. (B) DNA-binding domain (DBD) family composition of the TREND library. Of the 1,068 TFs, 729 were classified into 49 DBD families according to the Lambert et al. taxonomy (695 by direct symbol match and 34 resolved via HGNC-approved gene synonyms. Please see Table S1 for the full alias-to-canonical mapping); the 27 families with ≥4 members are shown individually, with the remaining 22 smaller families and 15 unclassified TFs grouped as ‘Other’. The six largest families (homeodomain, 176; C2H2 zinc finger, 122; basic helix-loop-helix, 63; nuclear receptor, 49; basic leucine zipper, 47; and forkhead, 32) together account for 67% of classified targets. Other (22 families + 15 unclassified) groups families with fewer than four members together with 15 confirmed TFs whose DBD family could not be assigned in the Lambert census. The remaining 339 targets include 248 proteins not present in the Lambert census (derived from protein-DNA interaction screens) and 91 entries classified as non-TFs (75 unlikely to be sequence-specific TFs and 16 ssDNA/RNA-binding proteins). (C) Total number of enhancer-reporter constructs (sensors) per DBD family. Each TF is represented by multiple sensors designed from distinct PPMs or PWMs, providing redundancy across motif variants. Total sensor count ranges from 80 (DM) to 11,283 (homeodomain), reflecting both the number of TFs per family and the diversity of available motif models per TF. (D) TREND library coverage of candidate lineage-specific cancer master transcription factors (MTFs) identified by the CaCTS algorithm across 34 TCGA tumor types^20^. Each bar represents one tumor type; red indicates MTFs present in the TREND library and grey indicates MTFs absent. Overall, 204 of 273 (74.7%) unique candidate MTFs are represented, with coverage ranging from 100% in uterine corpus endometrial carcinoma (UCEC), stomach adenocarcinoma (STAD), sarcoma (SARC), and lung squamous cell carcinoma (LUSC) to 58% in pheochromocytoma and paraganglioma (PCPG). Percentages above bars indicate the fraction of MTFs covered per tumor type. Full TCGA tumor type abbreviations follow standard nomenclature (https://gdc.cancer.gov/resources-tcga-users/tcga-code-tables/tcga-study-abbreviations). (E) TREND library coverage of core cell identity transcription factors across 15 anatomical systems, based on the top 10 specificity-ranked TFs per tissue from 233 cell and tissue types^21^. Tissues were grouped into 15 anatomical systems using an in-house rule-based classifier (full tissue→system mapping provided in Table S2. Overall, 354 of 503 (70.4%) unique core identity TFs are represented. Coverage is highest for the urogenital (83%) and respiratory (81%) systems and lowest for the endocrine system (55%). Percentages above bars indicate the fraction of unique core identity TFs covered per system. (F) Differential activity between ovarian cancer (OV8) and normal ovarian epithelial (IOSE) cells, highlighting cancer-selective synthetic enhancers (red) with high cancer-to-normal ratios. (G) Heatmap showing the relationship between motif variant rank within the PPM and enhancer specificity across TFs. Highly cancer-specific enhancers can arise from any motif rank, whether consensus or lower-ranked variants, rather than clustering at a particular rank. Ranks shown reflect retained variants after low-coverage filtering and are re-indexed contiguously.

Because TF binding motifs exhibit substantial diversity and context-dependent affinities^22,23^, we reasoned that individual motif variants could encode distinct binding strengths and activation thresholds. For therapeutic gene regulation, such threshold differences are critical, as disease specificity often depends not on the mere presence of TF activity, but on whether TF activity exceeds a level that distinguishes pathological from normal cellular states. We therefore hypothesized that systematically incorporating multiple motif variants per TF would generate enhancers spanning a range of activation thresholds, increasing the likelihood of identifying regulatory elements that enable precise, context-specific gene control.

To capture this range of thresholds, we performed Monte Carlo sampling on each PPM or PWM, generating 10,000 sequences per matrix by independent sampling from the position-wise nucleotide frequency distributions. The ten most frequently drawn unique sequences were retained per matrix: the top-ranked sequence as the consensus, and the remaining nine as affinity-modulating variants. Each enhancer was built as tandem repeats of a single TFBS sequence (consensus or variant) upstream of an adenoviral minimal promoter driving EGFP expression, linked to a unique DNA barcode co-transcribed with the reporter^17,24^. This design allows each enhancer’s activity to be quantified from barcode-linked RNA and DNA read counts. By pooling thousands of these constructs, we generated a library of 57,715 synthetic enhancer-reporter elements for high-throughput screening.

To demonstrate the utility of TREND for identifying cancer-specific regulatory elements, we applied the platform to ovarian cancer as a representative model system. The library was packaged into lentiviruses and transduced into human immortalized ovarian surface epithelial cells (IOSE), human ovarian cancer OVCAR-8 (OV8) cells, and murine ovarian cancer (ID8) cells. IOSE cells were used as a surrogate for normal ovarian surface epithelium because they retain key epithelial features and transcriptional similarity to primary ovarian surface epithelial cells while supporting efficient lentiviral transduction and high-coverage screening^25^.

For parallel quantification of enhancer activity, we used a barcode-based readout rather than FACS-based sorting. After delivering the pooled library to target cells, genomic DNA and total RNA were extracted. Construct-linked barcodes were quantified in both RNA and DNA by targeted next-generation sequencing (NGS) (Figure 1A); enhancer activity was calculated as the RNA/DNA barcode ratio.

Each enhancer was linked to multiple unique barcodes, enabling independent activity measurements and mitigating variability arising from lentiviral integration site effects, transcriptional noise, and enhancer-barcode mismatches. Analysis of the plasmid library confirmed robust barcode representation, with 77% of enhancers associated with more than 10 barcodes and a median of 30 barcodes per enhancer (Figure S1A and S1B). Enhancer activity was computed at the individual-barcode level and summarized as the median across associated barcodes within each biological replicate, followed by averaging across replicates to obtain a final per-construct activity score. Sequencing data were processed through a standardized pipeline including demultiplexing, UMI collapsing to correct for PCR amplification bias, and alignment to the reference library, with TPM normalization applied to account for differences in sequencing depth. Low-confidence elements were excluded based on DNA abundance and barcode representation thresholds (Figure S1C).

To evaluate the impact of filtering parameters on data quality and library coverage, we systematically varied reporting thresholds and barcode count cutoffs. Because every construct carries an identical minimal promoter with baseline transcriptional activity, barcodes with no detectable RNA signal are more likely to reflect insufficient cellular delivery or limited sequencing depth rather than true enhancer inactivity. We therefore defined a reporting threshold as the minimum fraction of enhancer-barcodes required to produce non-zero RNA counts at a given DNA abundance. For each reporting threshold, a corresponding minimum DNA copy number required for reliable detection can be inferred, and barcodes below this level were excluded from downstream analysis. Increasing the reporting threshold led to improved reproducibility between biological replicates (Figure S1D), consistent with the removal of low-abundance and intrinsically noisy barcodes. However, higher thresholds also reduced the number of retained enhancers (Figure S1E), indicating a trade-off between data quality and library coverage. Similarly, increasing the minimum barcode count per enhancer improved reproducibility (Figure S1F), while progressively reducing the number of enhancers that could be reliably quantified (Figure S1G). Based on these analyses, we selected a reporting threshold of 75% and a minimum barcode cutoff of ≥3, balancing reproducibility with library coverage. To assess cumulative attrition, we tracked enhancer retention across each stage of the pipeline, from initial design to post-filtering analysis. The majority of constructs were preserved through cloning and post-screening sequencing, with subsequent reductions primarily driven by reporting and barcode thresholds (Figure S1H). Notably, the ID8 dataset exhibited greater post-filtering attrition, likely reflecting lower transduction efficiency and reduced library coverage in this cell type compared with OV8 and IOSE cells.

Consistent with the broad and diverse library design, enhancer activity in IOSE and OV8 cells spanned a wide dynamic range, with most elements exhibiting comparable output across contexts and a distinct subset displaying markedly elevated OV8/IOSE activity ratios, identifying candidate cancer-selective synthetic enhancers (highlighted in red, Figure 1F). Downsampling analyses showed that replicate-to-replicate correlation was stable across sequencing depths (Figure S1I), indicating that activity measurements are robust to moderate reductions in coverage. However, lower depth reduced the number of reliably detected enhancers (Figure S1J). Activity profiles were also highly reproducible across biological replicates (Figure S2). These results indicate that while the quantitative measurements are stable, sufficient sequencing depth is required to maximize library coverage and ensure comprehensive enhancer identification.

Because transcription factor expression is often used as a proxy for regulatory activity, we next asked whether TF mRNA abundance alone could predict TREND-measured enhancer performance. Comparing enhancer activity with RNA-seq-based TF differential expression between OV8 and IOSE cells revealed no meaningful correlation (Figure S3A): the most cancer-selective enhancers were not associated with TFs showing the largest expression differences (Figure S3B), and conversely, enhancers linked to highly differentially expressed TFs did not exhibit strong cancer specificity (Figure S3C). Together, these results demonstrate that TF expression alone is a poor predictor of functional, context-specific enhancer activity, underscoring the value of direct functional screening for discovering context-specific regulatory elements.

Among the top cancer-specific enhancers, specificity did not correlate with motif variant rank within the PPM or PWM. For a given motif matrix, the most cancer-selective enhancer could derive from either the consensus sequence or lower-probability variants (Figure 1G), indicating that motif probability alone is not predictive of enhancer specificity. Instead, motif variants can encode distinct binding affinities that impose different activation thresholds. Sampling a large number of such variants enables identification of enhancers whose activation thresholds fall between TF activity levels in normal and cancer cells, so they remain silent in normal cells while becoming robustly activated in tumors. This threshold positioning is consistent with the sharp on/off behavior and high specificity observed for the selected enhancers. Accordingly, systematic incorporation of motif variants expands the accessible regulatory space and increases the likelihood of identifying synthetic enhancers with highly context-specific gene regulation. Consistent with the broad dynamic range observed in the screen, TREND revealed multiple classes of synthetic enhancers, including those selectively active in human cancer cells (high OV8/IOSE), mouse cancer cells (high ID8/IOSE), and shared cancer-selective elements active in both human and mouse cancer cells (high OV8/IOSE and ID8/IOSE), as well as enhancers preferentially active in immortalized human epithelial cells (high IOSE/OV8, high IOSE/ID8, or both). These results highlight the versatility of this approach in identifying diverse, context-dependent enhancer activities across species and cellular states (Figure S3D).

Collectively, these findings demonstrate that TREND enables direct functional assessment of synthetic enhancers, uncovering context-specific elements that are not predictable from TF expression or motif information alone.

### TREND identifies synthetic enhancers with robust cancer selectivity and TF-dependent activity

To test whether TREND-identified enhancers function as robust and cancer-selective regulatory elements, we individually cloned top candidates and evaluated their activity in OV8 and IOSE cells. Each enhancer was placed upstream of an adenoviral minimal promoter driving EGFP and delivered by lentivirus. Several motif variants from the MYC and E2F families consistently showed strong activity in OV8 while remaining largely inactive in IOSE (Figure 2A). Given their strong cancer selectivity and sequence diversity, MYC_v4 and E2F_v2 were selected as representative candidates: both produced strong, uniform EGFP fluorescence in OV8 with only background signal in IOSE (Figure 2B). Together, these results demonstrate that TREND can nominate synthetic enhancers with robust and cancer-selective activity.

**Figure 2.**
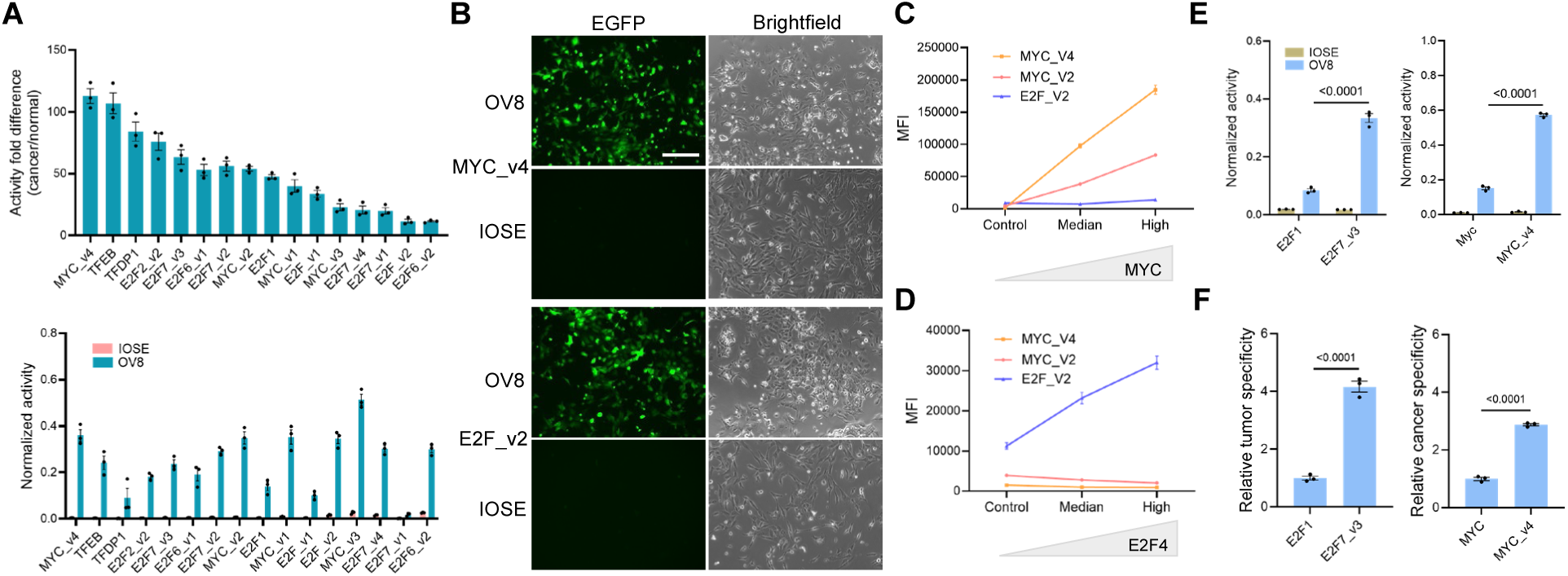
Functional validation of cancer-selective synthetic enhancers identified by TREND. (A) Quantitative reporter assays of individually cloned synthetic enhancers in human ovarian cancer (OV8) and normal ovarian epithelial (IOSE) cells measured by flow cytometry. Enhancer activity was normalized to the constitutive ubiquitin C (UbC) promoter control, and the activity fold difference was calculated as the ratio of normalized activity in OV8 cells relative to IOSE cells. v1, v2, v3, v4, etc. represent distinct motif variants derived from the same TF. (B) Representative fluorescence microscopy images of OV8 and IOSE cells transduced with MYC_v4 or E2F_v2 synthetic enhancers driving EGFP expression. Scale bar, 200 μm. (C and D) Reporter assays showing doxycycline dose-dependent activation of MYC_v4 and MYC_v2 (C) or E2F_v2 (D) enhancers following induction of their cognate TFs (MYC or E2F4) in IOSE cells. Data show median fluorescence intensity (MFI) of EGFP (n = 3) measured by flow cytometry. (E and F) Comparison of TREND-derived synthetic enhancers with previously reported designs in OV8 cells. E2F7_v3 and MYC_v4 were identified by TREND, whereas E2F1 and MYC represent previously reported enhancers. (E) Reporter activity driven by each enhancer. (F) Relative cancer specificity calculated as the ratio of enhancer activity in OV8 versus IOSE cells, with values normalized to the previously reported E2F1 and Myc enhancers. Data are mean ± s.e.m. from three independent experiments. Statistical significance was determined by unpaired two-tailed Student’s t-tests.

We next asked whether TREND-identified synthetic enhancers are functionally responsive to their intended TFs. To test this, we focused on two validated candidates, MYC_v4 and E2F_v2, and examined whether enhancer activity scaled with cognate TF abundance. Each enhancer was co-transduced with an inducible TF overexpression cassette, allowing TF abundance to be tuned by varying the inducer dose. Reporter assays in IOSE cells showed a clear dose-response: increasing MYC or E2F4 levels enhanced activity from the matched enhancer, whereas activity from the unmatched enhancer remained largely unchanged (Figure 2C, 2D, and S4). While this experiment focused on two representative enhancers rather than the entire library, the observed dose-dependent increase in activity with cognate TF abundance is consistent with the TF-centric design of the enhancer library. Together, these results support the expectation that TREND-nominated elements can reflect the activity of their corresponding TFs.

We previously showed that synthetic enhancers can achieve higher specificity than native cancer-specific promoters^7^. To further evaluate whether TREND-derived synthetic enhancers provide improved functional performance over those earlier manually designed constructs, we compared the top-performing TREND candidates with our prior design. Reporter assays showed that TREND-derived synthetic enhancers drove stronger activity in cancer cells while maintaining markedly improved specificity (Figure 2E and 2F). These results demonstrate the advantage of systematic, high-throughput functional screening over rational design for identifying enhancers with superior activity and specificity.

### Combinatorial TF sensing enhances cancer-restricted gene expression

Although individual TREND-derived enhancers exhibited high cancer selectivity, low-level background activity could still be detected in certain normal cells. To tighten control, we reasoned that endogenous TF activity, used here as a functional regulatory input, can be logically integrated rather than acted on individually. Gating transcription on defined combinations of independent TF activities (AND-gate logic) restricts expression to disease-specific transcriptional states that are unlikely to arise in healthy tissues.

To implement this logic, each TREND-derived enhancer was used to drive expression of one half of a split transcription factor: GAL4 DNA-binding domain (GAL4BD) fused to a leucine-zipper domain and VP16 activation domain (VP16AD) fused to its complementary zipper partner. Only when both enhancers are active are the two components produced simultaneously, allowing the zipper domains to associate and reconstitute a functional GAL4BD-VP16AD transcriptional activator (GAD). To identify effective leucine-zipper pairs for reconstitution, we assembled a library of 55 previously characterized heterodimeric coiled-coil domains, including the canonical SYNZIP set and additional validated interaction pairs from published studies ^26,27^ (Figure 3A). All possible 55 × 55 combinations were tested in a pooled format, with each candidate fused to either the GAL4BD (GAL4BD-zipper) or the VP16AD (zipper-VP16AD). Compatible pairings reconstitute a functional GAD, which activates a GAL4 promoter driving further expression of the same pair. This positive-feedback loop amplifies the transcripts of interacting pairs, so their relative abundance reports interaction strength. Because HEK293T cells do not express GAD endogenously, we incorporated a Tet-ON inducible system to transiently provide baseline GAD activity across all cells^28^: interacting pairs sustain themselves through the feedback loop after the pulse, while non-interacting pairs decay.

**Figure 3.**
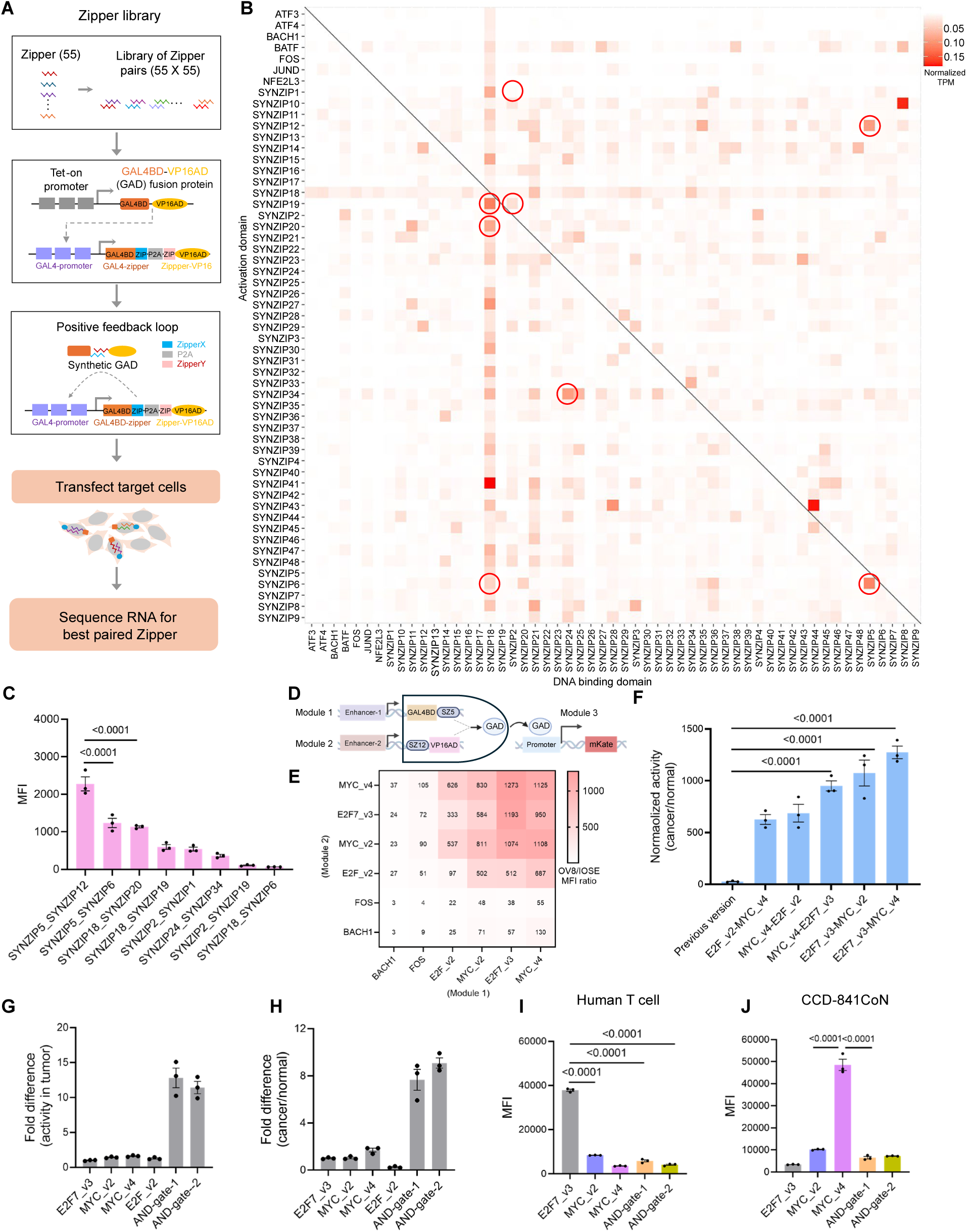
Enhancer-driven AND-gate circuits integrate multiple TF activities via protein-protein interactions. (A) Overview of the coiled-coil interaction screen used to identify optimal zipper pairs for circuit design. Each candidate coiled-coil domain was fused to either the GAL4BD-zipper or the zipper-VP16AD and expressed in HEK293T cells. Productive interactions between complementary zippers reconstitute a functional GAD, which drives reporter expression from a GAL4-responsive promoter. Transcript quantification by RNA sequencing was used to identify the most efficient interacting pairs. (B) Heatmap of pairwise interactions among 55 coiled-coil domain combinations, highlighting high-activity pairs (circled). (C) Flow-cytometry analysis of reporter activity for selected zipper pairs in HEK293T cells. (D) Schematic of the enhancer-driven AND-gate circuit, in which two distinct TREND enhancers independently drive GAL4BD-zipper and zipper-VP16AD expression. Productive coiled-coil pairing between the two proteins reconstitutes an active GAD complex, enforcing AND logic at the transcriptional level. (E) Cancer-to-normal activity ratios of AND-gate constructs assembled from synthetic enhancers MYC_v4, E2F7_v3, MYC_v2, E2F_v2, FOS, and BACH1, showing varying degrees of cancer selectivity. Heatmap values represent the mean from three independent experiments. (F) Comparison of cancer specificity between the top five AND-gate designs and a previously published RNA-based AND-gate configuration. Fold change represents the ratio of MFI in OV8 relative to IOSE. (G) Quantification of transcriptional output from AND-gate constructs relative to single synthetic enhancers. AND-gate architecture amplifies output strength, with activity normalized to the E2F7_v3 single-enhancer group. (H) Comparison of cancer-to-normal discrimination ratios between AND-gate and single-enhancer constructs. (I and J) Reporter activity of single synthetic enhancers and AND-gate constructs in human T cells and normal colon epithelial CCD-841 CoN cells. AND-gate-1 contains E2F7_v3 and MYC_v2 synthetic enhancers, and AND-gate-2 contains E2F7_v3 and MYC_v4 synthetic enhancers (E and F). Data are mean ± s.e.m. from three independent experiments. Statistical significance was determined by one-way ANOVA with Tukey’s multiple-comparisons test.

Viral delivery was performed at low multiplicity of infection so that each cell expressed only one candidate pair. RNA sequencing of zipper transcripts revealed a range of interaction strengths (Figure 3B). We selected representative pairs spanning weak to strong interactions (circled in Figure 3B) for downstream AND-gate construction. Flow cytometry showed that SYNZIP5-SYNZIP12 produced the strongest reporter activation (Figure 3C). To confirm AND-gate behavior, we tested whether partial configurations could activate the reporter. HEK293T cells were transiently transfected with two CMV-driven input modules (GAL4BD-SYNZIP2 and SYNZIP1-VP16AD) and a GAD-responsive output module (mKate). Reporter activity was tested under three input states: [1,0] (only module 1 present), [0,1] (only module 2 present), and [1,1] (both modules present). Reporter expression was detected only when both inputs were present (Figure S5A). The split-TF design therefore enforces AND-gate behavior.

To determine whether the same design remains functional when delivered by lentivirus for stable genomic integration, as intended for application in tumor models. Lentiviral delivery into human OV8 and murine BPPNM ovarian cancer cells produced robust activation in both lines (Figure S5B), confirming reliable AND-gate function in integrated form across species. Output strength was tunable by SYNZIP pair choice, demonstrating circuit adjustability. We then optimized SYNZIP positioning within the split TF, evaluating three fusion configurations. Activation occurred only when the SYNZIP was fused to the C-terminus of GAL4BD and the N-terminus of VP16AD (Figure S5C, S5D); alternative orientations failed to restore activity. Adding a nuclear localization signal (NLS) to VP16AD further increased reporter expression (Figure S5E).

We next asked whether TREND-derived enhancers could drive the AND-gate inputs. Six validated cancer-specific enhancers (MYC_v4, E2F7_v3, MYC_v2, E2F_v2, FOS, BACH1) were paired in all 6 × 6 two-input combinations. The modular circuit (Figure 3D) uses one enhancer to drive GAL4BD-SYNZIP5 (module 1) and the other to drive SYNZIP12-VP16AD (module 2); GAD is reconstituted, and the mKate reporter induced, only when both enhancers fire. The matrix screen identified several combinations with strong, cancer-restricted activity in OV8 and minimal activity in IOSE (Figure 3E, S6A, S6B). Specific enhancer pairs therefore cooperate through the AND-gate to enforce cancer-selective activation. The current circuit achieved substantially higher specificity than our earlier RNA-based AND-gate design^7^ (Figure 3F), and amplified transcriptional output relative to single-enhancer configurations (Figure 3G, 3H).

We further tested whether AND-gate integration could suppress the residual background activity seen with single enhancers in selected normal cells. In our prior study, MYC- and E2F-based enhancers showed minimal basal activity in primary human cells (dermal, ovarian, and ovarian microvascular endothelial), non-tumorigenic breast lines (MCF-10A, MCF-12A), and iPSCs^7^. However, MYC-based enhancers showed modest activity in a non-tumorigenic colonic epithelial line (CCD-841 CoN), and E2F-based enhancers in primary human T cells. In both contexts, AND-gate circuits sharply reduced background activity relative to single-enhancer constructs (Figure 3I, 3J). AND-gate integration therefore acts as a computational filter on TF-driven transcription, raising specificity while preserving or amplifying output. TREND-derived enhancers can thus be logically composed for high-fidelity transcriptional control.

### Tumor-restricted AND-gate circuits drive conditional combinatorial immunotherapy in vivo

Having shown that TREND-derived synthetic enhancers can be logically composed to integrate multiple TF activities, we next asked whether such transcriptional computation could be translated into tumor-restricted therapeutic programs in vivo: specifically, the conditional in situ production of multiple immunomodulatory payloads within tumors.

To translate this framework into murine in vivo models, we first applied the pipeline to ID8 ovarian cancer cells with IOSE as the normal reference (Figure S7A, S7B). When the genetically defined BPPNM model became available^29^, we prioritized it for downstream studies because it more closely reflects the genomic alterations of human ovarian cancer than ID8. To achieve broad coverage across model systems, we combined both human- and mouse-validated synthetic enhancers and evaluated their activity in BPPNM cells to identify cancer-selective candidates. We further evaluated these enhancers in two additional murine ovarian cancer models, KPCA and PPNM, which, together with BPPNM, constitute a panel of genetically engineered models encompassing the most frequent driver mutations observed in human ovarian cancer. From these analyses, several enhancer variants derived from the PPMs of ZBTB33 and MYC families showed the strongest and most consistent cancer-specific activity in BPPNM cells (Figure S7C-S7E). For AND-gate proof-of-concept, we selected two sequence-divergent variants, ZBTB33_v2 and MYC_v2.

We first assessed in vivo targeting precision and off-target expression, as these parameters are critical for clinical translation. BPPNM tumor-bearing mice and tumor-free C57BL/6 controls received intraperitoneal lentivirus encoding luciferase under the synthetic enhancers (Figure 4A). Bioluminescence was confined to tumor-bearing mice, with only background activity in healthy controls (Figure 4B, 4C). To rule out insufficient viral delivery as the cause of the silent normal-tissue signal, a CMV promoter control was included; this drove widespread expression in spleen, peritoneal wall, and tumor (Figure 4D, 4E). By contrast, both single-enhancer and AND-gate constructs restricted luciferase to tumor sites, with the AND-gate producing substantially stronger output than single enhancers. The AND-gate circuit therefore achieves robust, tumor-selective activation with minimal off-target expression in vivo.

**Figure 4.**
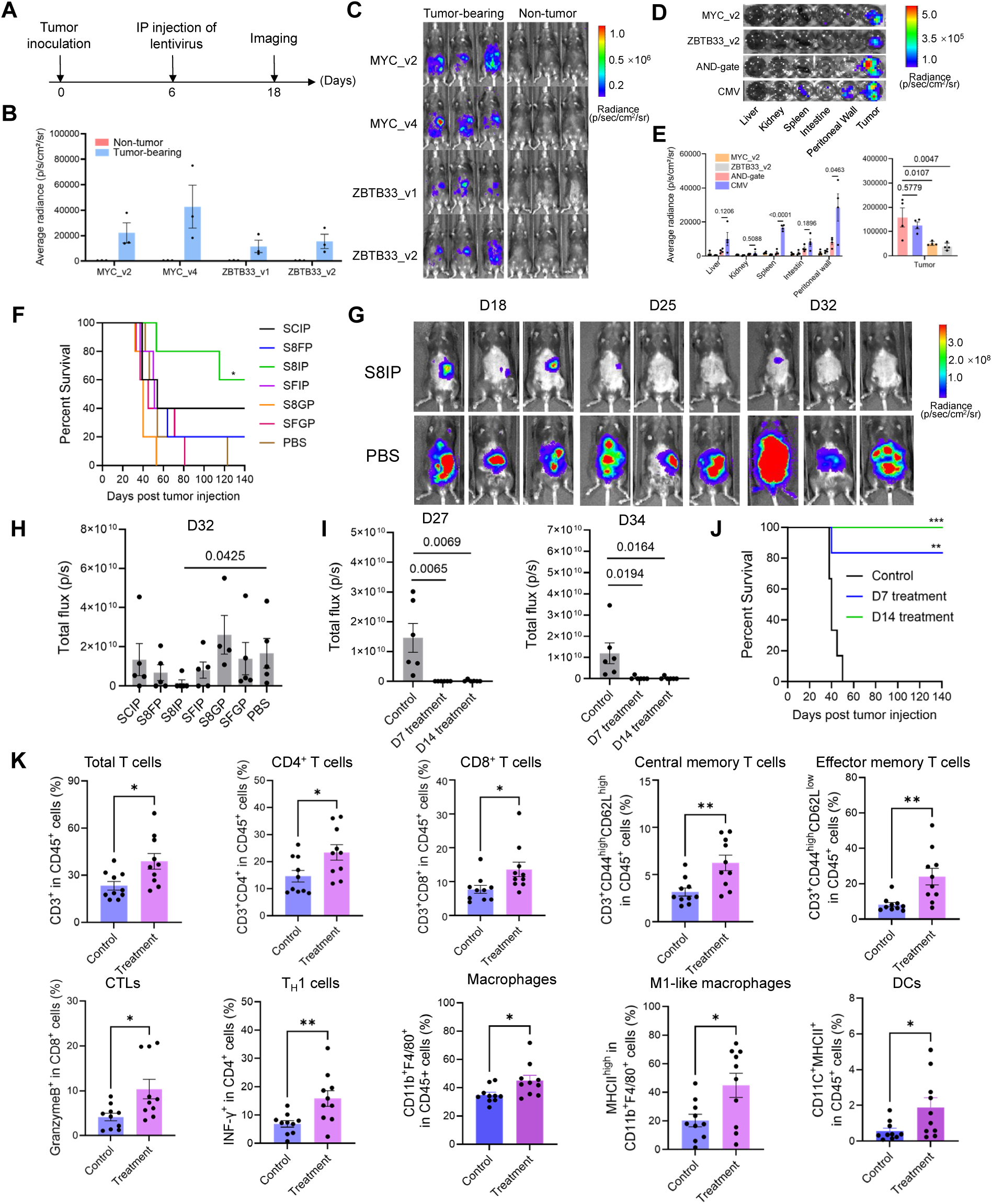
Tumor-restricted AND-gate circuits reprogram the tumor microenvironment and elicit durable therapeutic responses in murine ovarian cancer. (A) Experimental design for testing tumor specificity of individual TREND-derived synthetic enhancers in BPPNM tumor-bearing mice. (B) Quantification of bioluminescence signals from tumor-bearing and tumor-free mice (n = 3 mice per group). (C) Representative bioluminescence images of mice bearing BPPNM tumors or tumor-free controls following transduction with luciferase driven by single synthetic enhancers. (D) Ex vivo bioluminescence imaging of organs from mice transduced with single synthetic enhancers, enhancer-driven AND-gate circuits, or CMV promoter-driven luciferase. (E) Quantification of luciferase activity across major normal organs following delivery of single synthetic enhancer construct, AND-gate circuits or CMV enhancer constructs (n = 4 mice per group). (F) Survival curves of BPPNM tumor-bearing mice treated with AND-gate circuits encoding different combinations of immune effectors (n = 5 per group). (G) Representative bioluminescence images of tumor burden in treated mice. (H) Quantification of tumor growth on day 32 based on bioluminescence imaging (n = 5 per group). (I) Quantification of tumor growth in BPPNM tumor-bearing mice treated with a two-fold higher dose (relative to panels (F and H)) of the S8IP circuit, administered on day 7 or day 14 after tumor inoculation (n = 6 per group). (J) Survival curves of mice treated with the S8IP circuit on day 7 or 14, showing prolonged survival with both early and delayed dosing (n = 6 per group). (K) Flow-cytometry analysis of peritoneal immune cells following S8IP circuit treatment. BPPNM tumor-bearing mice were treated with the S8IP circuit at the dosage used in panels (I and J), administered on day 14 after tumor inoculation. Peritoneal wash was collected 7 days later for analysis. n = 10 per group. Statistical significance was determined by unpaired two-tailed t-tests, one-way ANOVA with Tukey’s multiple-comparisons test or log-rank (Mantel-Cox) test. *P < 0.05, **P < 0.01, ***P < 0.001 and ****P < 0.0001.

With tumor-restricted expression demonstrated in vivo, we next deployed this logic computation to orchestrate multiple immunomodulatory functions within tumors. We retained the same circuit architecture illustrated in Figure 3D, but reconfigured the output module to drive expression of therapeutic effectors rather than fluorescent reporters (Figure S8). Effective antitumor immunity is often constrained by multiple, concurrent barriers within the tumor microenvironment, including limited tumor antigen engagement, insufficient T-cell co-stimulation, and suppressive signaling that dampens effector responses^30^. To address these challenges in a coordinated manner, we assembled a panel of immunomodulators designed to target complementary aspects of tumor immunity. A surface T-cell engager (STE), consisting of a tumor-displayed anti-CD3ε single-chain variable fragment (scFv), was incorporated to promote direct T cell-tumor cell engagement, while CD80 provided co-stimulatory signaling to enhance T-cell activation^31–33^. IL-12 was incorporated to support T_H_1 polarization and effector differentiation, and Fms-related receptor tyrosine kinase 3 ligand (FLT3LG) was selected to expand dendritic cells (DCs) and promote antigen presentation^34,35^. To facilitate immune cell recruitment to the tumor milieu, we included CCL21 ^36^. In addition, the N-terminal fragment of Gasdermin D (GSDMD-N) was used to induce immunogenic tumor cell death, and an anti-PD-1 scFv was designed to locally alleviate checkpoint-mediated inhibition and restore T-cell function^37,38^.

We evaluated multiple combinations of these payloads in the BPPNM ovarian cancer model, including SCIP (STE, CCL21, IL-12, anti-PD-1 scFv), S8FP (STE, CD80, FLT3LG, anti-PD-1 scFv), S8IP (STE, CD80, IL-12, anti-PD-1 scFv), SFIP (STE, FLT3LG, IL-12, anti-PD-1 scFv), S8GP (STE, CD80, GSDMD-N, anti-PD-1 scFv), and SFGP (STE, FLT3LG, GSDMD-N, anti-PD-1 scFv), to identify therapeutic modules capable of reducing tumor burden and reprogramming the tumor microenvironment. Mice receiving the S8IP combination exhibited prolonged survival compared with controls (Figure 4F). Bioluminescence imaging confirmed reduced tumor burden, and quantification identified the S8IP module comprising STE, CD80, IL-12, and anti-PD-1 scFv as producing the strongest therapeutic effect (Figure 4G and 4H). Increasing the dosage of this circuit treatment further enhanced tumor control and extended survival, consistent with a dose-dependent therapeutic response (Figure 4I and 4J). Together, these results demonstrate that tumor-specific synthetic circuits coordinating multiple immunomodulators can elicit robust therapeutic benefit in vivo.

With potent, tumor-restricted activity established in vivo, we next examined whether circuit-driven execution of this combinatorial program reshapes the immune microenvironment. Flow cytometry analysis of peritoneal immune cells revealed broad remodeling of immune composition following S8IP treatment (Figure 4K). The proportions of total CD4⁺ and CD8⁺ T cells, central and effector memory T cell subsets, Granzyme B⁺ cytotoxic CD8⁺ T cells, and IFN-γ⁺ T_H_1 cells were markedly increased, consistent with enhanced adaptive immune activation. In parallel, increased frequencies of total macrophages, M1-like macrophages, and DCs suggest a shift in the myeloid compartment toward an immunostimulatory phenotype^39–41^. Together, these findings demonstrate that S8IP treatment not only reduces tumor burden but also reshapes the tumor microenvironment toward a coordinated pro-inflammatory state that supports durable antitumor immunity.

Additional immune subsets showed more selective changes. Both CD4⁺ and CD8⁺ T cells showed increased PD-1⁺ fractions, a marker of recent activation as well as checkpoint regulation (Figure S9A); given the expanded effector populations and inflammatory signature, this is more consistent with active engagement than terminal exhaustion. Other populations (NK, NKT, cytotoxic NK, T_reg_, Ly6C^high^ monocytes, and neutrophils) were largely unchanged (Figure S9B). Together, these findings suggest that the therapeutic circuits act primarily through enhancing T cell-mediated immunity, with potential contributions from myeloid effector activity, consistent with the design of the S8IP module to potentiate T cell-driven anti-tumor immune responses.

Collectively, these results demonstrate that TREND-derived synthetic enhancers, when integrated into AND-gate gene circuits, drive potent and tumor-restricted combination immunotherapy in vivo. By coupling transcriptional computation with combinatorial immunotherapy, this framework supports potent, spatially confined antitumor responses while minimizing off-target immune activation.

### TREND platform identifies activation-responsive synthetic enhancers for T cell activation-dependent gene control

To determine whether the TREND platform can be generalized beyond tumor targeting, we next applied it to T-cell engineering. In adoptive cell therapies such as chimeric antigen receptor (CAR) T-cell, coupling transgene expression to T-cell activation allows therapeutic payloads (cytokines or immune modulators) to be produced only on antigen engagement, enhancing efficacy while limiting systemic toxicity^6,42^. Activation-responsive promoters based on transcription factor binding motifs, particularly multimerized NFAT-responsive elements, have been widely used for this purpose. However, these promoters often exhibit basal activity in resting T cells, leading to unwanted background expression that can trigger premature effector release or systemic immune activation^43,44^. Scalable approaches for identifying activation-responsive enhancers with strong inducibility and tight off-state control are therefore needed to improve the safety and tunability of engineered T cell therapies.

To address this need, we applied TREND to identify synthetic enhancers that respond selectively and stringently to T cell activation. Because primary human T cells vary substantially in proliferation, transduction efficiency, and activation sensitivity across donors^45,46^, we sought to discover robust enhancers that perform reproducibly across donors rather than in any single donor. Primary T cells from two independent donors were activated, transduced with the pooled TREND library. After returning to a resting state, cells were stimulated, and enhancer-driven reporter activity was quantified by NGS of barcode-linked transcripts (Figure 5A). This workflow enabled direct comparison of enhancer activity between resting and activated states. From this screen, we identified a set of candidate activation-responsive enhancers, including NFKB1_v1, FOSL2, BCL, REL, RELA, FOSL1_v1, NFKB2, FOSL1_v2, NFKB1_v2, AP1_v3, AP1_v1, NFAT_v1, AP1_v2, and NFAT_v2, which showed strong induction upon T cell activation while remaining relatively silent in the resting state.

**Figure 5.**
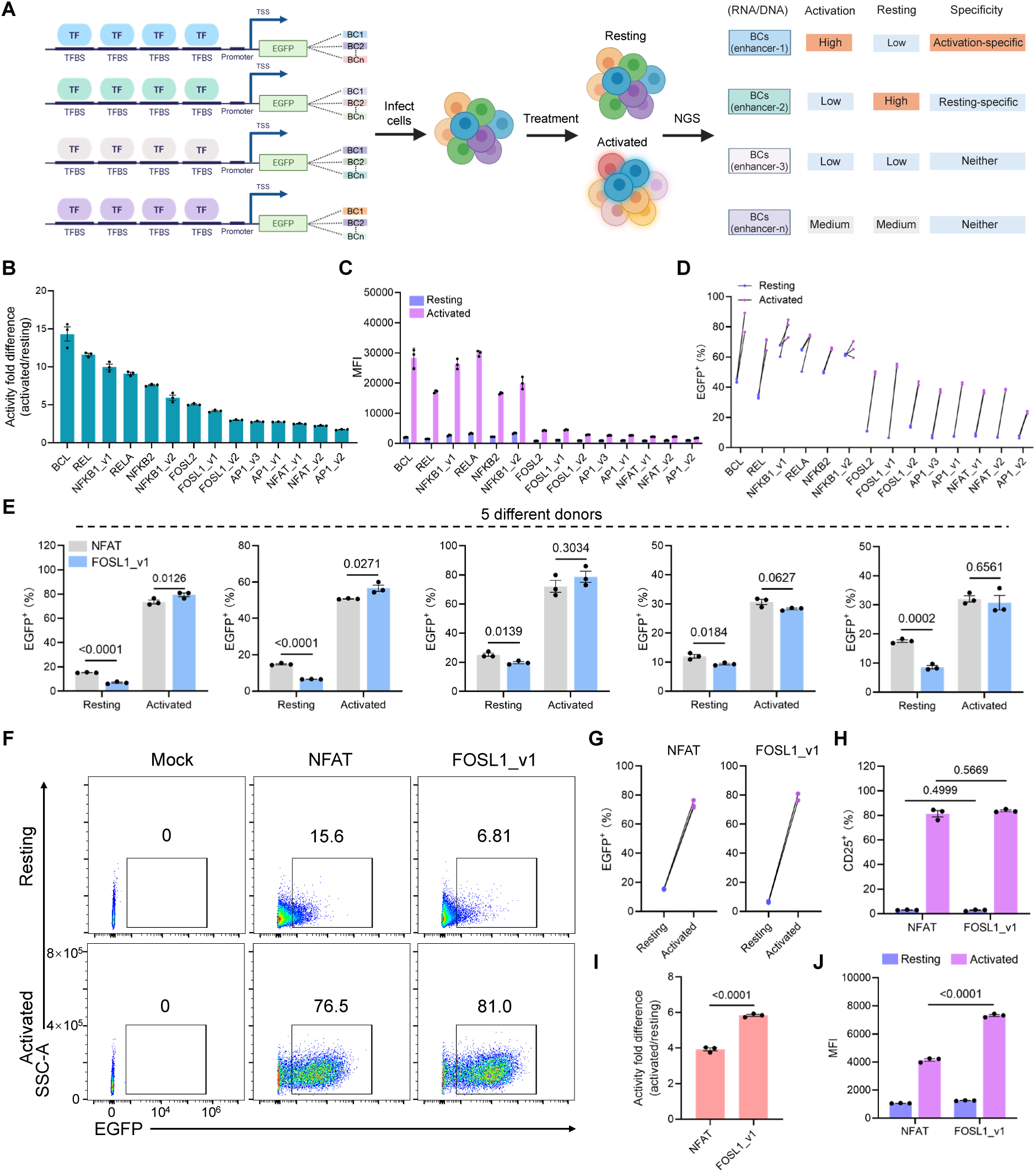
Identification of activation-responsive synthetic enhancers in T cells using TREND. (A) Schematic of the pooled screening workflow in human primary T cells. Cells were transduced with the TREND library, stimulated with anti-CD3/CD28 antibody, and enhancer activity was quantified by sequencing barcode-linked transcripts. (B) Fold difference in reporter activity (activated relative to resting condition) for all tested enhancers, identifying candidates responsive to T cell activation. (C) MFI of EGFP expression driven by individual enhancers in activated versus resting T cells. (D) Percentage of EGFP^+^ T cells under activated and resting conditions, showing activation-induced enhancer responsiveness. (E) Comparison of EGFP^+^ cell percentages for the canonical NFAT-motif-based enhancer and the TREND-derived FOSL1_v1 enhancer under activated and resting conditions. (F) Representative flow cytometry analysis showing EGFP expression driven by NFAT and FOSL1_v1 enhancers in resting and activated T cells. (G) Quantification of EGFP⁺ cell percentages from panel (F). (H) Proportion of CD25^+^ T cells transduced with NFAT or FOSL1_v1 enhancer under resting and activated conditions. (I) Fold difference in reporter activity (activated relative to resting condition) for NFAT and FOSL1_v1 enhancers. (J) MFI of EGFP expression for NFAT and FOSL1_v1 enhancers in activated and resting T cells. Data are mean ± s.e.m. from three independent experiments. Statistical significance was determined by unpaired two-tailed t-tests.

We next validated these candidates individually in primary human T cells from three independent donors. Across donors, most enhancers exhibited markedly higher activity in the activated state than in the resting state (Figure 5B, 5C, and S10A-S10F). Notably, several candidates, including NFKB1_v1, BCL, REL, RELA, and NFKB1_v2, also showed substantially elevated basal EGFP expression in resting T cells, indicating incomplete off-state silencing (Figure 5D, S10C and S10F). Among all elements tested, FOSL1_v1 consistently demonstrated the highest activation specificity, driving robust reporter expression upon stimulation while maintaining near-background activity in resting cells across all donors.

To confirm that these results reflected enhancer-intrinsic regulatory performance rather than differences in T cell activation, we measured expression of CD25, a canonical activation marker, and observed comparable induction across all conditions (Figure S10G). This supports the interpretation that variation in enhancer-driven reporter output arose from differences in regulatory performance rather than unequal activation efficiency. Consistent with this interpretation, direct comparison with a conventional NFAT-motif-based enhancer showed that FOSL1_v1 exhibited markedly lower basal expression in resting T cells while driving stronger reporter induction upon stimulation (Figure 5E-5J and S11). T cells transduced with either construct displayed similar CD25 expression, further supporting equivalent activation states (Figure 5H). Collectively, these results demonstrate that FOSL1_v1 functions as a tighter and more precise transcriptional switch than existing NFAT-based designs, potentially enabling more stringent activation-dependent control of therapeutic payload expression in engineered T cells.

Taken together, these findings demonstrate that TREND can be extended beyond tumor-targeting contexts to discover activation-responsive synthetic enhancers in primary human T cells, underscoring its versatility as a platform for discovering context-specific regulatory elements.

## Discussion

Synthetic enhancers that enable precise, context-dependent gene expression are increasingly important for both fundamental biology and therapeutic engineering ^47^. In this study, we developed TREND, a TF-guided screening framework that systematically identifies synthetic enhancers with high cellular specificity and context-dependent activity. The platform provides a broadly applicable approach for precise gene control across diverse biological systems. Existing approaches for enhancer discovery often depend on cell-type-specific regulatory datasets, such as ATAC-seq, DNase-seq, or ChIP-seq-derived enhancer maps, or on computational models trained on those datasets ^10,11,15–17^. While these methods can reliably recover enhancer activity in well-characterized contexts, they typically require new reference data or model retraining when applied to different cell types or disease states, which limits their flexibility.

Because TF activities vary across cell types and physiological states, assaying a broad library of TF-responsive sequences enables the native transcriptional environment to reveal which motifs are preferentially active in a given context. The TREND library comprises 57,715 enhancer designs targeting 1,068 proteins with annotated DNA-binding motifs, of which 729 are confirmed sequence-specific transcription factors representing 49 DNA-binding domain families. The library captures 75% of candidate lineage-specific cancer MTFs across 34 TCGA tumor types and 70% of core tissue identity TFs across 233 cell and tissue types, providing dense coverage of the regulatory factors most relevant to cell identity and disease. Each construct is built from tandem repeats of a single motif sequence derived from the PPM or PWM of that factor; when screened, each serves as a functional reporter of endogenous transcriptional programs, linking cellular state to enhancer performance. By assaying motif-driven transcriptional activity directly in native cellular environments, TREND identifies synthetic enhancers with cell-type or state specificity, without relying on pre-annotated regulatory libraries or computational models.

Compared with prior motif-based synthetic enhancer screening strategies^17^, TREND introduces both conceptual and technical advances that expand the scope and utility of TF-guided regulatory design. Our earlier SPECS platform screened a limited panel of TF motifs using FACS-based enrichment. TREND scales this approach by systematically sampling over 1,000 DNA-binding factors with multiple motif variants per factor and replaces FACS enrichment with barcode-linked RNA/DNA quantification, giving a direct quantitative measure of activity for each construct. Graded motif variants per TF further enable systematic exploration of enhancer design space, facilitating identification of elements with optimized activity and specificity. Unlike conventional MPRAs, which evaluate genomic regulatory regions, TREND interrogates TF motif space using a fully synthetic, modular design, enabling context-independent exploration of regulatory logic. Unlike ML-based design, which requires high-quality epigenomic training data per cell type, TREND is empirical and applies to contexts where training data are limited or unavailable. Finally, TREND’s scalable architecture supports direct integration with downstream gene circuit engineering. A summary comparison of these approaches is provided in Table 1.

**Table 1:** Comparison of TREND with prior synthetic enhancer screening approaches.

| **Feature** | **SPECS (2019)** | **MPRA (typical)** | **TREND** |
| --- | --- | --- | --- |
| Library source | Motif subset | Genomic regions | TF motif space |
| TF coverage | Limited | Indirect | 1,068 targets |
| Variant design | Consensus only | None | Systematic variants |
| Readout | FACS<br>enrichment | Barcode<br>RNA/DNA | Barcode<br>RNA/DNA |
| Quantitative | Semi | Yes | Yes |
| Compatible with fragile cells | No | Yes | Yes |
| Requires prior datasets | No | Yes | No |

Applying TREND to human and murine ovarian cancer systems uncovered synthetic enhancers that robustly distinguished cancer cells from non-tumorigenic epithelial cells. Candidates from the MYC and E2F families displayed strong cancer-selective activity and responded quantitatively to cognate TF abundance, suggesting that TREND-identified enhancers can serve as compact, modular reporters of disease-associated transcriptional programs. Compared with earlier manually designed synthetic enhancers^7^, the TREND-derived elements exhibited higher output and improved cancer selectivity.

Although a single TREND-derived enhancer often achieved strong specificity, low-level activity could occasionally be detected in non-malignant cells. To further tighten control, we implemented a protein-interaction-based AND-gate circuit that integrates signals from two enhancers: each enhancer drives one half of a split transcription factor (GAL4BD-SYNZIP and SYNZIP-VP16AD), and only when both enhancers are active do the two halves interact through their coiled-coil SYNZIP domains to reconstitute a functional GAD. This configuration suppresses background when one enhancer exhibits leakage while maintaining strong on-target expression when both enhancers are active. More broadly, our results support a general principle, consistent with prior demonstrations of combinatorial specificity in synthetic circuits, that signal integration through logic-gated architectures enhances regulatory precision while preserving inducible expression^48,49^.

We next established a proof-of-concept for in vivo application using these enhancer-driven AND-gate circuits in the BPPNM murine ovarian cancer model. A critical distinction from our prior xenograft work is that the BPPNM model is a genetically defined syngeneic system that recapitulates the most prevalent mutations or alterations in human high-grade serous ovarian cancer (Brca1, Trp53, Pten, Nf1, and Myc)^29^. Because this model retains a fully intact immune system, it enables rigorous characterization of the immune response elicited by circuit-driven immunotherapy, an aspect that could not be adequately assessed in immunodeficient xenograft settings. In this model, cancer-specific enhancer pairs drove effector modules encoding complementary immune mediators. These circuits restricted activity to tumor sites, reduced tumor burden, and improved survival while promoting a pro-inflammatory tumor microenvironment. Among tested configurations, the S8IP module (STE, CD80, IL-12, anti-PD-1 scFv) produced the strongest dose-dependent therapeutic response. That circuit-driven immunomodulation reshaped the tumor immune landscape within an immunocompetent host underscores translational relevance and distinguishes this work from xenograft-based demonstrations. Future work could focus on identifying the minimal synergistic effector combinations within S8IP to reduce circuit complexity, and on integrating multiple modules into one or two vectors to simplify manufacturing and tighten control over component stoichiometry.

Extending TREND beyond tumor-targeting, we applied it to primary human T cells to uncover synthetic enhancers that precisely couple transgene expression to T cell activation. The resulting constructs exhibited greater activation-dependent expression and lower basal activity than the conventional NFAT-motif-based promoter used in earlier cytokine-secreting CAR-T designs^42,43^. Among them, FOSL1_v1 reproducibly linked reporter expression to T cell stimulation across donors. Recent rational-engineering and computational-design efforts have produced improved activation-responsive promoters^6,50,51^; FOSL1_v1 complements these by showing that unbiased functional screening can identify regulatory elements with favorable properties without prior knowledge of the underlying logic. Although the difference in resting-state baseline expression between the NFAT-motif-based enhancer and FOSL1_v1 may appear modest (15.6% versus 6.81% EGFP⁺ cells), recent work has demonstrated that even this level of basal leakiness can drive excessive effector release and toxicity in vivo^6^. These observations underscore that small differences in resting-state activity can have outsized functional consequences in therapeutic settings, a consideration that may become increasingly important as inducible gene control systems move toward clinical translation.

Several aspects of TREND could be refined. First, the current library focuses on enhancers built from single TF motifs in tandem, which simplifies design but does not explore the cooperative or combinatorial motif architectures of natural regulatory elements^52,53^. Future studies integrating multiple motif types could provide additional tuning flexibility and may more closely recapitulate the logic of endogenous enhancers. In addition, incorporating machine-learning-guided optimization could help generate synthetic enhancers with improved performance beyond those currently represented in the library.

To facilitate adoption and reuse, we provide an open-source pipeline and interactive dashboard (https://github.com/SyntheticImmunity/TREND-Bioinformatics-Pipeline) that wraps the analyses presented here. Distributed as a single containerized image, the dashboard offers browsable views of the full ∼2.7 million construct library, one-click reproduction of the deposited activity tables, and a scaffolded path for applying TREND to new datasets. The entire pipeline runs on a standard laptop with no local bioinformatics infrastructure required.

Overall, TREND offers a versatile experimental framework for discovering synthetic enhancers with strong contextual specificity. By directly assaying motif-driven transcription across cellular contexts, the approach provides a scalable path for tailoring regulatory elements to specific cellular or disease settings, with potential to enable safer gene therapies, more precise cell-based immunotherapies, and a broader toolkit for programmable synthetic biology.

## Method

### TREND library construction

For the construction of the TREND library, we collected motif matrices for human transcription factors from two complementary sources. Position probability matrices (PPMs) were obtained from the Bioconductor MotifDb package (v1.46.0), which aggregates experimentally determined TF binding profiles from SELEX, HT-SELEX, protein-binding microarray (PBM), and ChIP-seq assays across major databases including JASPAR, UniPROBE, hPDI, ScerTF, Stamlab, and cisBP. Position weight matrices (PWMs) were obtained from the ENCODE motif compendium^54^, which provides complementary motif models inferred from large-scale ChIP-seq data across hundreds of TFs. All source matrices were standardized to PPM format prior to sequence generation: PWMs were converted to frequency space by exponentiation and per-column renormalization so that each column contained non-negative probabilities summing to one.

For each standardized matrix, we performed Monte Carlo sampling to identify representative binding sequences. Ten thousand independent sequences were generated per matrix by sampling one nucleotide at each position according to the position-wise probability distribution, treating positions as independent. The resulting sequences were ranked by draw frequency, and the ten most frequent unique sequences that passed design filters were retained per matrix. Design filters required that each retained sequence (i) had not already been selected from another matrix for the same TF, and (ii) contained no recognition site for the restriction enzymes used in downstream cloning (AgeI, AscI, BamHI, SbfI), both within the sequence itself and within its immediate flanking cloning context. The first qualifying sequence was designated the consensus; the next nine were retained as affinity-modulating variants. The consensus reflects the most frequent qualifying draw rather than the analytical PPM mode. All ten sequences are screened in parallel. For the large majority of matrices, this empirical consensus coincides with the analytical PPM mode. Sequence generation was implemented in Python 3 using numpy.random.choice for per-position nucleotide sampling from renormalized PPM columns and collections.Counter.most_common for frequency ranking of the resulting sequences.

Each synthetic enhancer design comprised tandem repeats of a single transcription factor binding site (TFBS) sequence (consensus or variant) separated by 3 bp spacers, appended iteratively until the total length of the variable region approached 121 bp, ensuring consistent construct size across designs. The complete pool of variable regions was synthesized as an array-based oligonucleotide library (oligopool) by Agilent Technologies, Inc. (Santa Clara, CA, USA) using array-based DNA synthesis. Each design in the pool contained a variable region comprising tandem TFBS repeats, flanked by constant elements including universal primer sites and restriction enzyme sites. Each oligonucleotide was then amplified by PCR using primers containing 20 bp random sequences, introducing a unique barcode for quantifying barcode abundance at both the DNA and RNA levels.

The barcode-containing fragments were first digested with AgeI and SbfI and cloned into a lentiviral backbone, generating an intermediate vector. This intermediate construct was then digested again with AscI and BamHI to insert the adenoviral minimal promoter and EGFP reporter gene, yielding the final expression cassette. In this configuration, each TFBS-based synthetic enhancer regulates EGFP expression, and its associated barcode is positioned at the 3’ end of the EGFP transcript, enabling quantification of barcode abundance at both the DNA and RNA levels. This design enables quantitative assessment of enhancer activity through barcode-linked expression analysis.

### SYNZIP library construction

The Zipper library was designed to systematically evaluate pairwise interactions among 55 synthetic coiled-coil “zipper” domains. All possible 55 × 55 zipper pairs were synthesized as an oligonucleotide pool in the format zipper₁-P2A-zipper₂, where the self-cleaving P2A peptide separates the two coding sequences. The oligonucleotide pool was synthesized by Agilent Technologies, Inc. (Santa Clara, CA, USA) using array-based DNA synthesis. The synthesized fragments were PCR-amplified and cloned into a GAL4-responsive expression vector containing the GAL4 promoter, such that each fragment was inserted between GAL4BD and VP16AD to generate the fusion construct GAL4BD-zipper₁-P2A-zipper₂-VP16AD. This configuration allows co-expression of paired zipper domains within the same cell under control of the GAL4 promoter, enabling high-throughput functional screening of zipper-zipper interactions.

### Cell culture and cell lines

The human ovarian carcinoma cell line OV8 was originally established at the National Cancer Institute (NCI) and obtained from existing laboratory stocks. BPPNM, KPCA, and PPNM cell lines were gifts from the laboratory of Robert A. Weinberg (Massachusetts Institute of Technology). The ID8 murine ovarian cancer cell line was a gift from the laboratory of Katherine F. Roby (University of Kansas Medical Center). Lenti-X HEK293T (Takara, 632180), CCD-841 CoN cells (American Type Culture Collection, CRL-1790), IOSE (Applied Biological Materials, T1074), and OVCAR8 were cultured in DMEM (Corning, 10-013-CV) supplemented with 10% fetal bovine serum (FBS; Corning, 35-010-CV), 1% non-essential amino acids (MEM/NEAA; Hyclone, 16777-186), and 1% penicillin–streptomycin (Life Technologies, 15140-122). ID8 cells were cultured in DMEM supplemented with 10% heat-inactivated FBS and 1% penicillin–streptomycin. BPPNM, PPNM, and KPCA cells were cultured in DMEM supplemented with 1% insulin–transferrin–selenium (Thermo Fisher Scientific, 41400045), 2 ng/mL EGF (Life Technologies, PMG8041), 4% heat-inactivated FBS, and 1% penicillin–streptomycin. All cells were maintained at 37 °C in a humidified atmosphere containing 5% CO_2_ and were routinely tested negative for mycoplasma contamination.

### Primary human T cells

Leukoreduction collars from anonymous healthy platelet donors were obtained through the Brigham and Women’s Hospital Crimson Core Laboratory. CD3^+^ T cells were isolated from the white blood cell fraction using the RosetteSep Human T Cell Enrichment Kit (STEMCELL Technologies, 15021), followed by density gradient centrifugation with Lymphocyte Separation Medium (Corning, 25-072-CV) and SepMate-50 tubes (STEMCELL Technologies, 85450).

### Lentivirus Production and Transduction

Lentiviral particles were generated by reverse transfection of HEK293T cells in a 24-well format using FuGENE-HD (Promega, E2311). All plasmid DNA was normalized to 0.1 µg/µl in sterile water. For each well, a plasmid mixture containing 0.2 µg of lentiviral transfer vector, 0.1 µg psPAX2 (Addgene #12260), and 0.1 µg pVSV-G (Addgene #138479) was prepared. In parallel, 2.4 µl of FuGENE-HD was diluted in 20 µl of Opti-MEM (Thermo Fisher Scientific, 31985070), mixed with the plasmid solution, vortexed briefly, and incubated at room temperature for 15-20 min to allow complex formation.

HEK293T cells were maintained below 80% confluency, and the culture medium was replaced with fresh DMEM 2-4 hours before transfection to promote optimal viral yield. Cells were trypsinized, counted, and resuspended in DMEM to a final concentration of 3.6 × 10⁶ cells/mL. A volume of 100 µl of the cell suspension (3.6 × 10^5^ cells) was added to each transfection tube containing the FuGENE-DNA complexes, mixed gently, and plated into wells pre-filled with 300 µl of fresh DMEM. Eighteen to twenty hours after transfection, the medium was replaced with 500 µl of fresh DMEM. Viral supernatants were harvested 44-48 hours post-transfection, clarified by centrifugation, and filtered through a 0.45 µm syringe filter (Pall Corporation, 4614). Filtered lentivirus was used immediately or stored at 4 °C for up to 3 days or -80 °C for long-term storage.

For transduction, 1.25 × 10^6^ target cells were infected with serial dilutions of filtered viral supernatant in the presence of 8 µg/mL polybrene (Sigma) overnight. Medium was replaced the following day. For comparative experiments, viral titers were normalized using the Lenti-X p24 Rapid Titer Kit (Takara, 631476) before use.

### Inducible transcription factor overexpression and reporter assay

A doxycycline-inducible (Tet-On) system was used to regulate TF expression in IOSE cells. IOSE cells were co-infected with three lentiviral vectors: one encoding the reverse tetracycline-controlled transactivator 3 (rtTA3) under the UbC promoter, a second encoding doxycycline-inducible MYC or E2F4 (Tet-On system), and a third carrying the synthetic enhancer-driven EGFP reporter (MYC_v4 or E2F_v2). Seventy-two hours after infection, cells were treated with increasing concentrations of doxycycline (0, 100, and 1000 ng/mL; Thermo Scientific Chemicals, J60579) for 72 hours to induce graded TF expression. Reporter activity was quantified by flow cytometry (Beckman Coulter, CytoFLEX), and median EGFP fluorescence intensity was analyzed using FlowJo software.

### T cell activation and transduction

Cryopreserved primary human T cells were thawed (day 0) and cultured in RPMI 1640 (Corning, 10-040-CV) supplemented with 10% heat-inactivated FBS, 10 mM HEPES (Life Technologies, 15630080), 0.1 mM NEAA, 1 mM sodium pyruvate (Life Technologies, 11360-070), 1% penicillin–streptomycin, 50 µM 2-mercaptoethanol (Sigma-Aldrich, M3148-25ML), and 50 IU/mL recombinant human IL-2 (NCI). T cells were stimulated using the Human CD3/CD28 T Cell Activator (STEMCELL Technologies, 10971) according to the manufacturer’s protocol. On day 2, activated T cells were resuspended in lentiviral supernatant and centrifuged at 2,000 × g for 2 h at 32 °C (spinoculation). Viral supernatants were then replaced with fresh culture medium. After transduction, T cells were maintained by diluting 1:2 daily with fresh medium to sustain exponential growth and viability.

For screening and validation of synthetic enhancers, T cells were transduced on day 2 and cultured as described above. On day 10, cells were restimulated with the Human CD3/CD28 T Cell Activator and collected 24 hours later for analysis.

### Lentiviral library transduction and sequencing

Pooled lentiviral libraries, including the TREND and Zipper constructs, were packaged and transduced (with an estimated MOI of 5 for TREND or 0.3 for Zipper constructs) into the respective target cell types using the standard lentiviral production and transduction workflow described above. To ensure reproducibility and minimize sampling bias, we maintained >100-fold representation of each library element throughout the screening pipeline. Five days post-transduction, genomic DNA and total RNA were isolated from pooled cell populations using the AllPrep DNA/RNA Mini Kit (Qiagen, 80204).

DNA library preparation: Barcoded genomic DNA fragments were first amplified by a limited-cycle PCR (3 cycles) using primers that introduced unique molecular identifiers (UMIs), sample indices, and Illumina adapter sequences, enabling simultaneous UMI tagging and adapter addition while minimizing amplification bias. The resulting products were then amplified in a second PCR (15 cycles) using universal Illumina primers to enrich adapter-ligated fragments and generate sequencing-ready libraries. Amplicons of the expected size (∼132 bp) were gel-purified, pooled, and cleaned prior to sequencing.

RNA library preparation: For each sample, 2 µg of total RNA was reverse-transcribed using primers containing a 5’ Illumina adapter, a unique molecular identifier (UMI), and a sample-specific index sequence. Reverse transcription was performed at 42 °C for 60 min, followed by heat inactivation at 85 °C for 5 min. After RNase A treatment, cDNA was amplified by PCR (∼15 cycles) using Illumina-compatible universal primers to generate sequencing-ready amplicons (∼132 bp). Where indicated, emulsion PCR was tested using Q5 polymerase supplemented with 0.02 mg/mL BSA to reduce amplification bias.

Sequencing libraries were quantified, pooled, and sequenced at the Dana-Farber Cancer Institute Molecular Biology Core Facility on an Illumina NovaSeq platform using 150 bp paired-end reads.

### Barcode quantification and enhancer activity analysis

Sequencing data were processed using a custom in-house pipeline implemented on the Harvard Medical School O2 high-performance computing cluster. Read orientation (“flipping”) and sample demultiplexing were performed using in-house Python scripts guided by predefined barcode–sample key tables. Demultiplexed reads were collapsed using FASTX-Toolkit (fastx_collapser) to remove duplicate reads, and the resulting files were aligned to the reference library using Bowtie2 with parameters optimized for short, high-similarity sequences. Barcode sequences were then extracted from read pairs using in-house Python and R scripts, and identical barcodes were aggregated to minimize PCR and sequencing biases. Aligned reads were processed with SAMtools to obtain barcode-level count tables for DNA and RNA libraries.

Downstream analysis was performed using an in-house R script. Barcodes with zero DNA or RNA counts were excluded, and constructs represented by fewer than a specified minimal number of independent barcodes were filtered out to ensure reproducibility. For each barcode, the RNA/DNA ratio was calculated as a quantitative measure of enhancer activity. Within each biological replicate, barcode-level ratios were summarized by their median value, and the mean of these replicate medians was used to represent the final enhancer activity score for each construct. To ensure quantitative robustness, we examined the distribution of DNA barcode for each sample and defined a lower DNA-abundance threshold such that at least 75% of constructs (cancer-specific enhancer project) or 50% of constructs (activation-specific enhancer project) remained transcriptionally active above that cutoff. Barcodes passing this threshold were aggregated by enhancer identity, and constructs meeting coverage criteria were retained for downstream analysis. These activity scores were then used for comparative analysis between tumor and control cell lines or between activated and resting T-cell states. For the cancer-specific enhancer validation experiments, top candidates were selected based on consistent performance across biological replicates. For the activation-specific enhancer validation, top candidates were prioritized from the full library screen in individual donors, with additional preference given to enhancers containing motifs derived from FOS, AP-1, or NFAT families.

### Bulk RNA-seq analysis

Total RNA from OV8 and IOSE cells was extracted and submitted to Azenta Life Sciences (GENEWIZ, South Plainfield, NJ, USA) for library preparation and sequencing. RNA quantity was measured using a Qubit 2.0 Fluorometer (Thermo Fisher Scientific), and RNA integrity was assessed with an Agilent TapeStation 4200 (Agilent Technologies). RNA-seq libraries were prepared using the NEBNext Ultra II RNA Library Prep Kit for Illumina (New England Biolabs) with poly(A) selection, following the manufacturer’s protocol. Briefly, mRNA was enriched using Oligo(dT) beads, fragmented at 94 °C for 15 min, and reverse-transcribed to generate first- and second-strand cDNA. The resulting cDNA fragments were end-repaired, adenylated at the 3’ ends, ligated to Illumina adapters, and amplified by limited-cycle PCR. Library size and concentration were verified using the Agilent TapeStation and Qubit fluorometer.

Sequencing was performed on an Illumina NovaSeq 6000 platform in a 2 × 150 bp paired-end configuration, targeting approximately 30 million reads per sample. Raw sequencing data were converted to FASTQ format and demultiplexed using bcl2fastq v2.20, allowing one mismatch for index recognition. Adapter trimming and quality filtering were performed with Trimmomatic v0.36. Clean reads were aligned to the human reference genome (ENSEMBL) using STAR v2.5.2b, and gene-level read counts were generated with featureCounts v1.5.2 from the Subread package, considering only uniquely mapped reads within annotated exon regions. Transcript abundance was normalized as TPM for downstream analysis.

### Animal experiments

All animal care and experiments were conducted at the Dana-Farber Cancer Institute Animal Research Facility under protocol 19-014, reviewed and approved by the Institutional Animal Care and Use Committee. Mice were housed in ventilated cages (up to five mice per cage) under standard conditions including controlled light/dark cycles, temperature, and humidity.

### In vivo antitumor efficacy validation

Three million BPPNM cells engineered to stably express firefly luciferase (BPPNM-Luc) were suspended in a 1:1 mixture of Matrigel (Corning, 47743-710) and phosphate-buffered saline (PBS) and injected intraperitoneally into 6-8-week-old female C57BL/6 mice (Jackson Laboratory, 000664). Tumor progression was monitored by bioluminescence imaging using an IVIS Spectrum imaging system (Xenogen), and images were analyzed with Living Image Software v4.7.4. On day 6 post-implantation, mice were imaged to assess baseline tumor burden; animals showing outlier radiance values were excluded to minimize bias. The remaining mice with comparable radiance levels were then randomly assigned to experimental groups.

For in vivo delivery of the S8IP circuit, lentiviruses encoding each circuit component (module 1, module 2, STE, CD80, IL12, and anti-PD1 antibody) were produced as described above and concentrated tenfold using a Sorvall WX 80+ ultracentrifuge. Concentrated lentiviruses were combined at a volume ratio of module 1 : module 2 : STE : CD80 : IL12 : anti-PD1 = 1 : 1 : 0.5 : 0.5 : 0.5 : 0.5. Lentiviruses for other circuit groups, including SCIP, S8FP, SFIP, S8GP, and SFGP were produced using the same method. Each mouse received an intraperitoneal injection of 400 μL of the viral mixture (∼1 × 10^10^ viral particles/mL), while the higher-dose group received 400 μL at approximately 2 × 10¹⁰ viral particles/mL. For control groups, lentiviruses were mixed at identical ratios, substituting the S8IP components with vectors encoding the output reporter rtTA3 or PBS. Viral titers were quantified and normalized before injection using the Lenti-X p24 Rapid Titer Kit (Takara, 631476).

### Flow cytometry for immune analysis

BPPNM tumor-bearing mice received an intraperitoneal injection of the S8IP circuit at a dose of 400 μL of viral mixture (∼2 × 10^10^ viral particles/mL) on day 14 after tumor inoculation. Peritoneal wash was collected 7 days later for analysis. Briefly, mice were euthanized and injected with 5 mL of cold PBS to collect the peritoneal fluid. The collected fluid was centrifuged at 800 x g for 5 min at 4 °C, and the cell pellet was subjected to red blood cell lysis with ACK buffer for 10 min at room temperature. After washing with PBS, cells were resuspended in FACS buffer and passed through 70-μm strainers prior to downstream applications.

For surface staining, cells were first incubated with TruStain FcX™ PLUS (BioLegend, 156603) for 10 min on ice, followed by staining with fluorophore-conjugated antibodies (1:200 dilution, 30 min at 4 °C). After washing with PBS, cells were incubated with Zombie NIR viability dye (BioLegend, 423105; 1:1000 in PBS) for 30 min at 4 °C.

Antibodies used for lymphoid cell analysis include BUV737-CD3 (BD Biosciences, 612821), BV510-CD45 (BioLegend, 103137), BV650-CD4 (BioLegend, 100545), BV711-CD8 (BioLegend, 100747), BV785-CD62L (BioLegend, 104440), Percp Cy5.5-TIM3 (BioLegend, 119717), PE-NK1.1 (BioLegend, 156503), PE/Dazzle 594-CD44 (BioLegend, 103055), PE-Cy7- CD19 (BioLegend, 115519), APC-PD1 (BioLegend, 135210), AF700-CD25 (BioLegend, 102024) and APC-Cy7-CD11b (BioLegend, 101225). APC-Cy7-CD11b was used to exclude CD11b^+^ cells from lymphoid analysis.

Antibodies used for myeloid cell analysis include BV421-Ly6G (BioLegend, 127627), BV510-CD45 (BioLegend, 103137), BV650-CD11c (BioLegend, 117339), BV785-CD11b (BioLegend, 101243), Percp Cy5.5-Ly6C (BioLegend, 128011), PE-XCR1 (BioLegend, 148204), PE/Cy5-F4/80 (BioLegend, 123111), PE-Cy7-CD172a (BioLegend, 144007), APC-CD206 (BioLegend, 141708), AF700-MHCⅡ (BioLegend, 107621), APC-Cy7-CD3 (BioLegend, 100221) and APC-Cy7-CD19 (BioLegend, 115529). APC-Cy7-CD3 and APC-Cy7-CD19 were used to exclude CD3^+^ cells and CD19^+^ cells from myeloid analysis.

For intracellular staining, cells were stimulated in culture at 1 × 10^6^ cells/mL with 2 μl/mL of activation cocktail and GolgiPlug (BD Biosciences, 550583) for 4 h at 37 °C. Cells were washed once and pre-incubated with TruStain FcX™ PLUS for 10 min on ice, followed by surface antibody staining (1:200, 30 min at 4 °C) and Zombie NIR staining (1:1000, 20 min in PBS). After PBS wash, cells were then fixed and permeabilized using the FoxP3/Transcription Factor Staining Buffer Set (eBioscience, 00-5523-00), washed twice with permeabilization buffer, and incubated with intracellular antibodies (1:100 in permeabilization buffer, 1 h at room temperature).

Antibodies used for intracellular cytokines analysis include BUV737-CD3 (BD Biosciences, 612821), BV421-granzyme B (BioLegend, 515409), BV510-CD45 (BioLegend, 103137), BV650-CD4 (BioLegend, 100545), BV711-CD8 (BioLegend, 100747), BV785-IFN-γ (BioLegend, 505838), PE-Perforin (ebioscience, 12-9392-82), PE-Cy7-NK1.1 (BioLegend, 156513), APC-IL-4 (BioLegend, 504105), AF700-Foxp3 (BioLegend, 126421) and APC-Cy7-CD11b (BioLegend, 101225).

Samples were acquired with Cytek Aurora using SpectroFlo v.3.1.0. Collected data were analysed using FlowJo v.10.8.1 Software.

### In vivo specificity validation

Three million BPPNM cells were resuspended in a 1:1 mixture of Matrigel (Corning, 47743-710) and PBS and injected intraperitoneally into 6–8-week-old female C57BL/6 mice (Jackson Laboratory). On day 6 post-tumor implantation, mice received 400 μL of lentivirus (∼2 × 10^10^ viral particles/mL) encoding firefly luciferase driven by synthetic enhancers. Luciferase activity was measured on day 18 by in vivo bioluminescence imaging using an IVIS Spectrum imaging system (Xenogen), and analyzed with Living Image Software v4.7.4.

For AND-gate and CMV control groups, lentiviruses were concentrated tenfold using a Sorvall WX 80+ ultracentrifuge and mixed at a volume ratio of module 1 : module 2 : luciferase = 1 : 1 : 2. On day 6, each mouse was administered 400 μL of the viral mixture (∼2 × 10^10^ viral particles/mL) or lentivirus encoding luciferase under the CMV promoter. On day 28, the liver, kidney, spleen, intestine, peritoneal wall, and tumor tissues were harvested for ex vivo bioluminescence imaging.

## Statistical analysis

Data were analyzed using GraphPad Prism v10.4.1. The statistical tests applied to each dataset are specified in the corresponding figure legends. Unless otherwise noted, values are presented as mean ± s.e.m. Statistical significance was assessed using unpaired two-tailed Student’s t-tests, log-rank (Mantel-Cox) test, one-way analysis of variance (ANOVA) or two-way ANOVA followed by Tukey’s multiple comparisons test. Statistical difference was considered significant at *P < 0.05, **P < 0.01, ***P < 0.001. **** P < 0.0001.

## Materials availability

The TREND library and key plasmids generated in this study will be deposited in Addgene upon publication. Other materials are available from the corresponding author upon reasonable request.

## Code and data availability

Processed sequencing data, including post-alignment RNA and DNA barcode count tables for all screens, and summarized screening results will be made available upon publication. The accompanying R scripts for enhancer activity quantification (barcode filtering, normalization, and activity scoring) will be made available alongside the processed data. Raw sequencing data are available from the corresponding author upon reasonable request, in accordance with institutional data-sharing policies.

## Acknowledgements

We thank the Molecular Biology Core Facility (MBCF) at Dana-Farber Cancer Institute for technical support and the Animal Research Facility (ARF) for expert animal care. This work was supported by the Cancer Research Institute (CRI) CLIP Award, the Department of Defense Ovarian Cancer Research Program (OC220238/W81XWH-22-OCRP-IIRA), and the Parker Institute for Cancer Immunotherapy.

## Author contributions

Y.J., C.-Y.C., and M.-R.W. conceived the study and developed the methodology. Y.J. and M.-R.W. analyzed and interpreted the data and wrote the manuscript. Y.J., C.-Y.C., B.Z., R.W., A.M.S., S.H., Y.-N.L., H.Y., T.S., Y.-C.W. and Z.W. performed experiments and collected data. L.S., Y.W., Y.W. and E.B.A. provided technical support and contributed to data interpretation. M.-R.W. supervised the study and coordinated the overall project.

## Competing interests

M.-R.W., L.S., C.-Y.C., B.Z., H.Y., S.H., Y.J., and R.W. are co-inventors on patent applications filed by Dana-Farber Cancer Institute related to the technologies described in this work, including international patent application WO2025029678A2 (PCT/US2024/039879) and U.S. Provisional Application No. 63/806,926, titled “Synthetic Immunomodulatory Nucleic Acids for Ovarian Cancer.” M.-R.W. and L.S. are co-founders of TroGen Therapeutics, a company developing gene circuit-based cancer therapies, and hold equity in the company. All other authors declare no competing interests.

---

**Figure S1.**
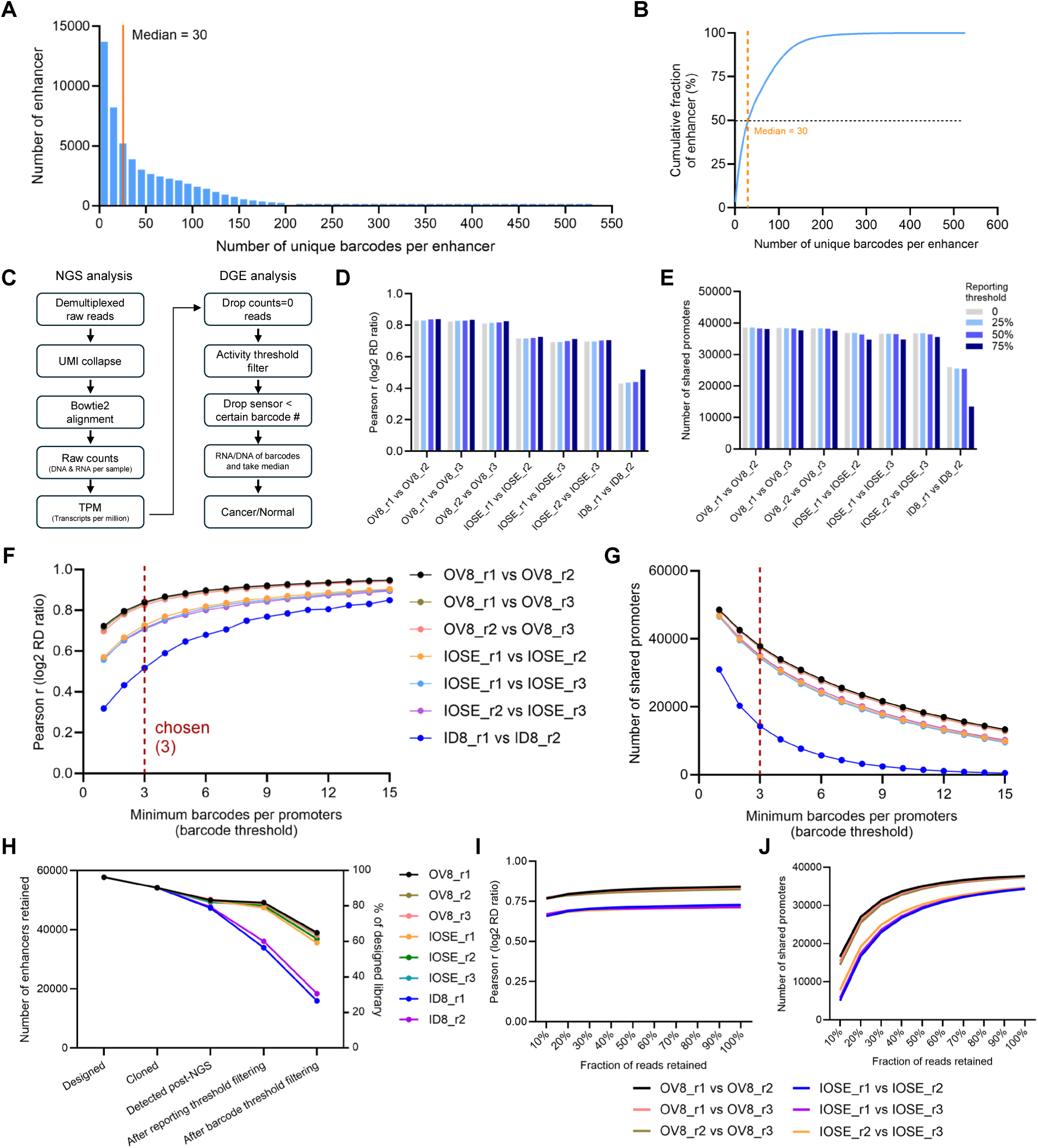
Quality control and filtering analyses for TREND library screening. (A) Histogram of unique barcode counts per enhancer in the plasmid library. The median of 30 barcodes per enhancer is indicated (orange line). (B) Cumulative distribution of the data shown in (A). The dashed horizontal line marks the 50th percentile; the dashed vertical line marks 30 barcodes per enhancer. (C) Schematic of the TREND data-processing pipeline. Left (NGS analysis): demultiplexing of raw reads, UMI collapsing, Bowtie2 alignment, per-sample raw DNA and RNA count assembly, and transcripts-per-million (TPM) normalization. Right (differential gene expression analysis): removal of zero-count entries, activity-threshold filtering, minimum-barcode filtering, per-barcode RNA/DNA ratio calculation with per-enhancer median aggregation, and downstream classification of enhancer activity. (D) Pearson correlation of log₂(RNA/DNA) values between pairs of biological replicates (OV8, IOSE, ID8) at four reporting thresholds (0%, 25%, 50%, 75%; color-coded). (E) Number of enhancers retained (shared promoters) across the same replicate pairs and reporting thresholds as in (D). (F) Pearson correlation of log₂(RNA/DNA) values between biological replicate pairs as a function of minimum barcode count per enhancer (barcode threshold, 1-15). The red dashed line at barcode threshold = 3 indicates the cutoff used in downstream analyses. (G) Number of enhancers retained across the same replicate pairs and barcode threshold range as in (F). (H) Per-sample enhancer retention across five stages of the pipeline: designed, cloned, detected post-NGS, after reporting-threshold filtering (75%), and after barcode-threshold filtering (≥3 barcodes per enhancer). Left y-axis: number of enhancers retained; right y-axis: percentage of designed enhancers retained. Each line represents one biological replicate (OV8, IOSE, ID8). (I) Downsampling analysis of Pearson correlation between biological replicate pairs across decreasing fractions of retained sequencing reads (10%-100%). (J) Number of enhancers retained across the same downsampling series as in (I).

**Figure S2.**
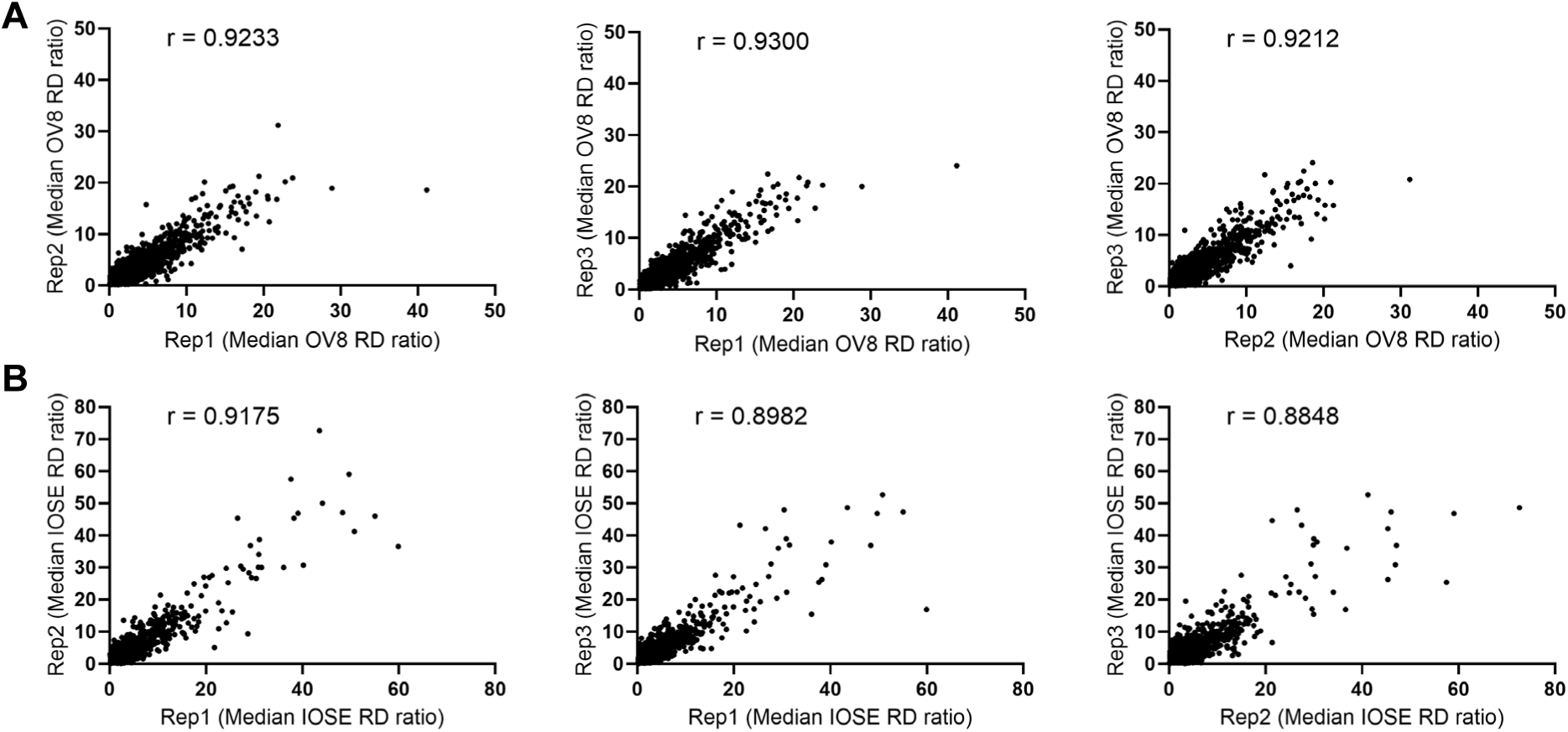
TREND screening shows high reproducibility across biological replicates. (A and B) Correlation of synthetic enhancer activity profiles between independent biological screening replicates in OV8 (A) and IOSE (B) cells. Each point represents a synthetic enhancer, with RNA output normalized to DNA input (RNA/DNA ratio, referred to as RD ratio). Pearson correlation coefficients indicate high reproducibility between TREND screening replicates.

**Figure S3.**
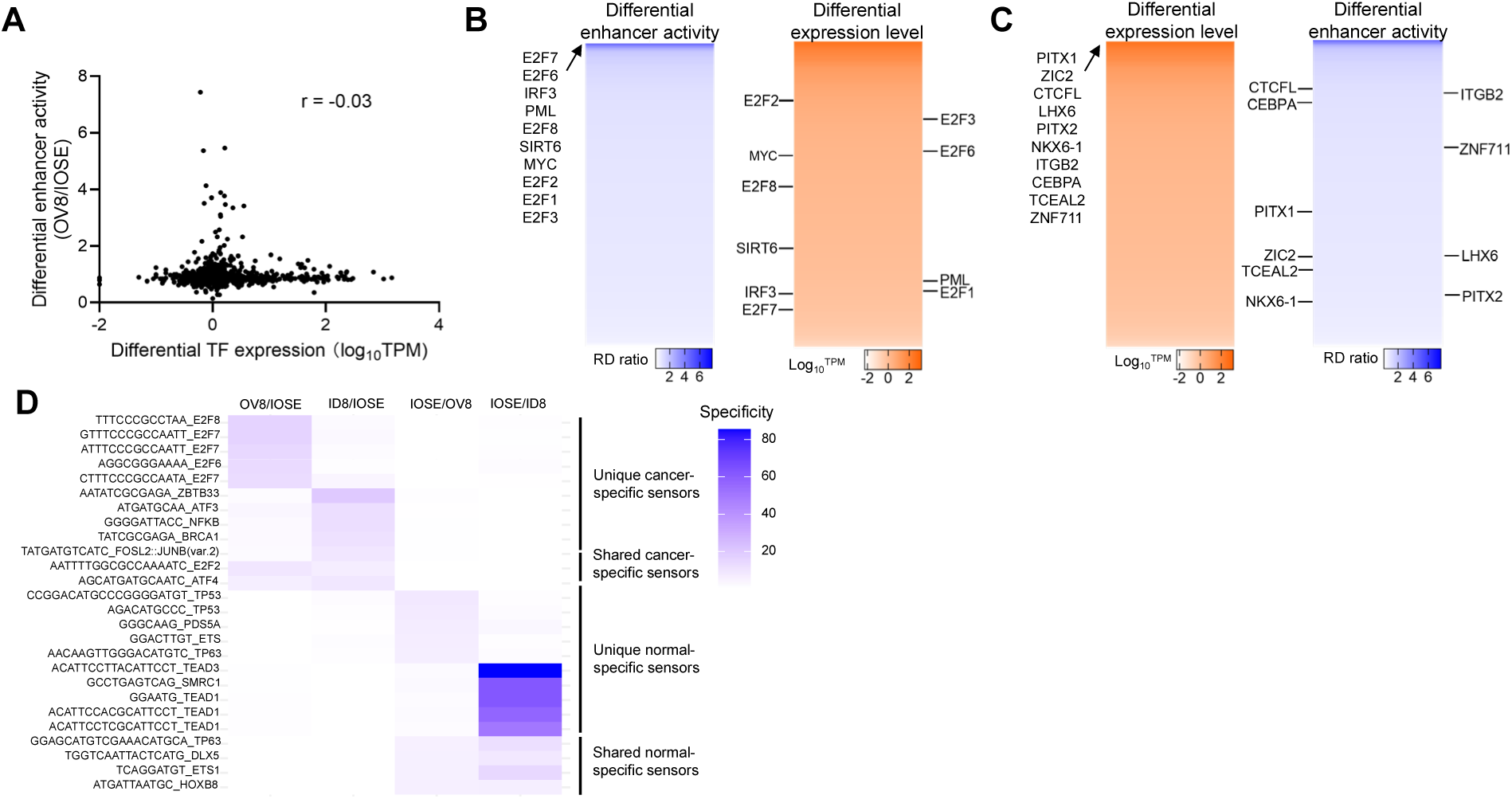
Relationship between TF expression and synthetic enhancer activity. (A) Scatterplot comparing differential TF mRNA expression (OV8 versus IOSE) with the corresponding differential activity (OV8 versus IOSE) of the synthetic enhancers responsive to each TF, measured by TREND. Each point represents a specific TF. (B and C) Heatmaps comparing differential enhancer activity (OV8/IOSE activity ratio) of TREND-identified synthetic enhancers with the corresponding TF differential expression (OV8/IOSE, RNA-seq). TFs associated with top-performing cancer-specific enhancers are dispersed throughout the TF differential-expression landscape rather than concentrated among highly differentially expressed TFs (B), and conversely, TFs with high transcript differences are distributed across a wide range of enhancer specificity without a consistent trend (C). (D) Representative classes of enhancers, including those selective for human cancer cells (high OV8/IOSE), mouse cancer cells (high ID8/IOSE), and shared cancer-selective elements active in both human and mouse cancer cells (high OV8/IOSE and ID8/IOSE), as well as enhancers preferentially active in normal human epithelial cells (high IOSE/OV8, high IOSE/ID8, or both). The heatmap values denote the specificity, defined as the activity ratio between cell types.

**Figure S4.**
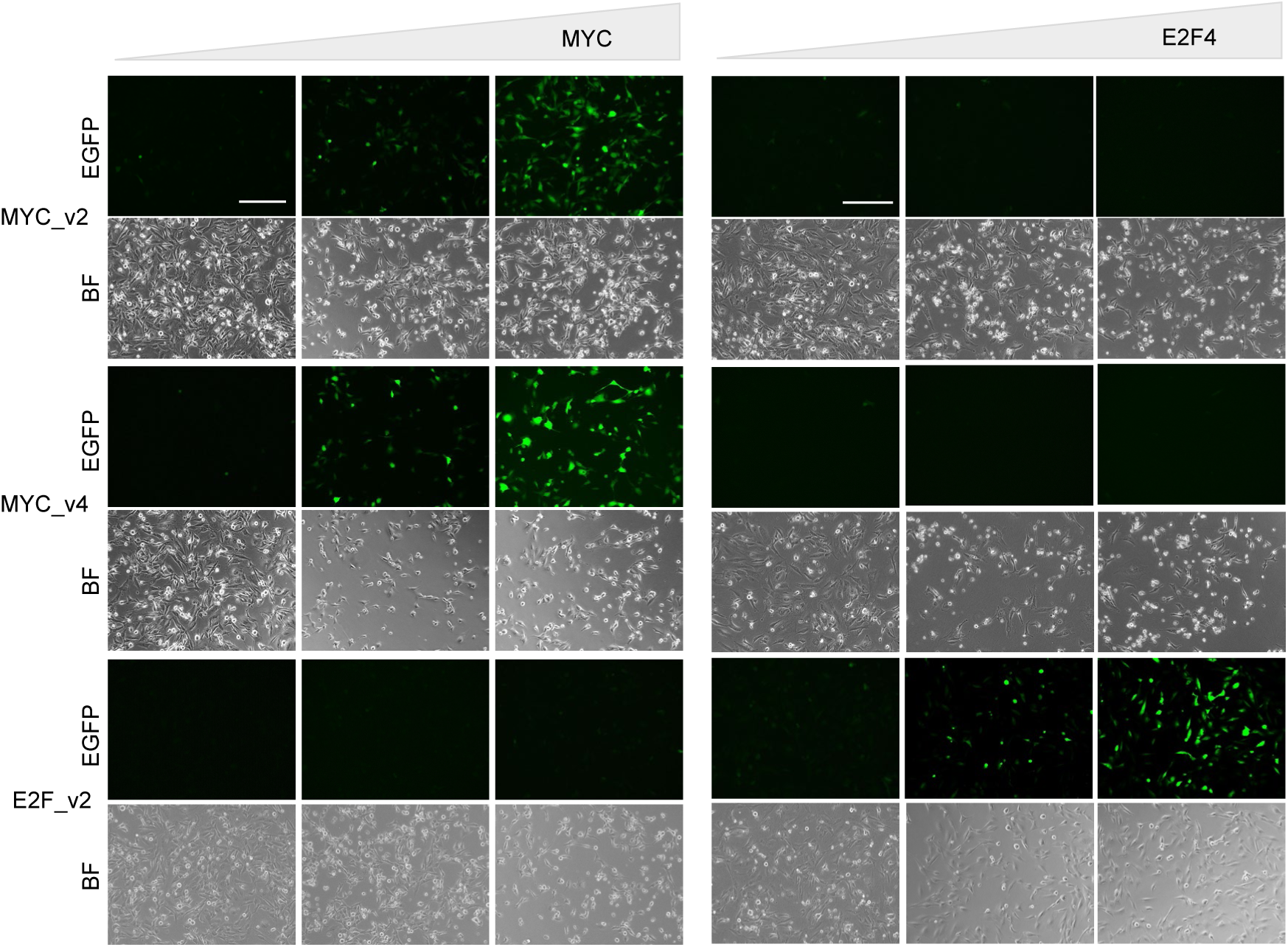
Dose-response of TREND-derived synthetic enhancers to TF abundance. Fluorescence microscopy images of IOSE cells co-transduced with MYC_v4 or E2F_v2 synthetic enhancers and a Tet-ON-inducible TF expression cassette for overexpressing MYC or E2F4. Images show graded reporter activation in response to TF abundance. Scale bar, 300 μm. Data are representative of three independent experiments.

**Figure S5.**
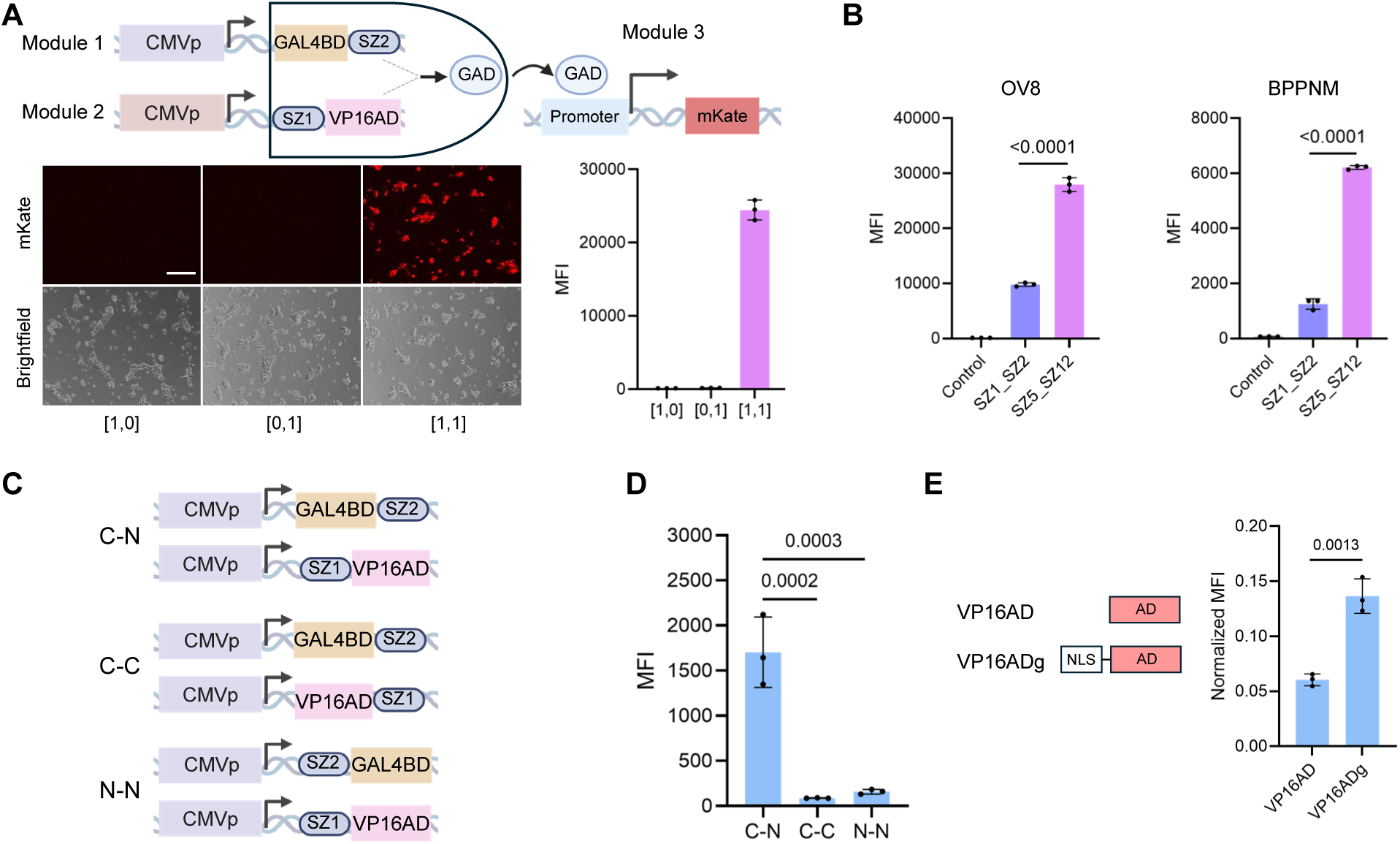
Orientation-dependent effects and nuclear localization enhance SYNZIP-based AND-gate performance. (A) Reporter assays of partial AND-gate configurations, showing that incomplete circuits containing only one enhancer input ([1,0] or [0,1]) fail to activate transcription. Scale bar, 200 μm. (B) Validation of SYNZIP-based AND-gate activity in human (OV8) and mouse (BPPNM) ovarian cancer cells. (C) Schematic illustrating different orientations of SYNZIP fusion to GAL4BD and VP16AD domains. (D) Quantification of reporter activity across different SYNZIP fusion orientations. (E) Effect of a nuclear localization signal (NLS) to VP16AD on downstream reporter expression. Data represent mean ± s.e.m. from three independent experiments. Statistical significance was determined by unpaired two-tailed t-tests or one-way ANOVA with Tukey’s multiple-comparisons test.

**Figure S6.**
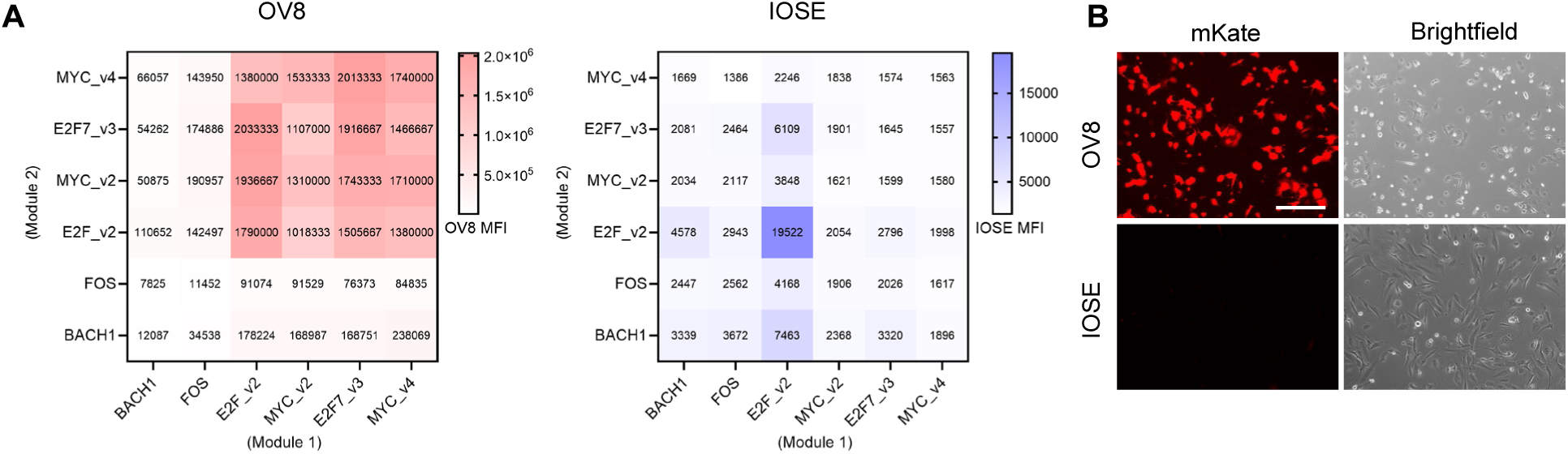
Functional evaluation of AND-gate circuits constructed from TREND-derived synthetic enhancers. (A) Reporter assays comparing the activity of AND-gate constructs in OV8 and IOSE cells. Each construct was assembled from validated TREND-derived enhancers (MYC_v4, E2F7_v3, MYC_v2, E2F_v2, FOS, and BACH1), evaluated in different pairwise combinations. Heatmap values represent the mean from three independent experiments. (B) Representative fluorescence-microscopy images of mKate reporter in OV8 and IOSE cells transduced with the AND-gate construct.

**Figure S7.**
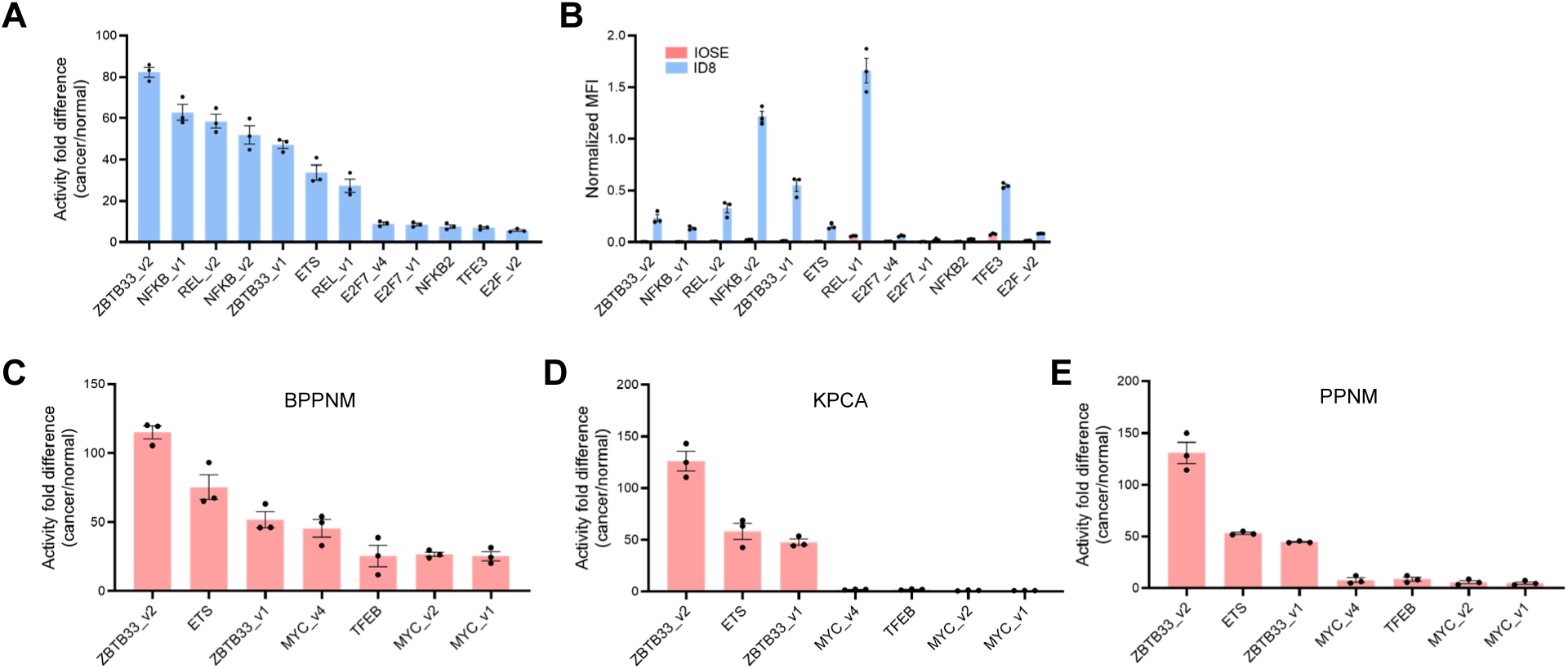
Screening and validation of murine cancer-specific synthetic enhancers. (A) Tumor specificity of TREND-derived synthetic enhancers in ID8 mouse ovarian cancer cells, quantified as the fold difference in activity relative to normal human ovarian epithelial cells (ID8/IOSE). (B) Normalized enhancer activity in ID8 and IOSE cells corresponding to the data in panel a, illustrating relative enhancer activity across cancer and normal contexts. MFI values were normalized to the UbC promoter control. (C to E) Validation of synthetic enhancer activity across additional mouse ovarian cancer cell lines, including BPPNM (C), KPCA (D), and PPNM (E). Activity fold difference represents the ratio of MFI in BPPNM/IOSE, KPCA/IOSE, and PPNM/IOSE, respectively. Data are mean ± s.e.m. from three independent experiments.

**Figure S8.**
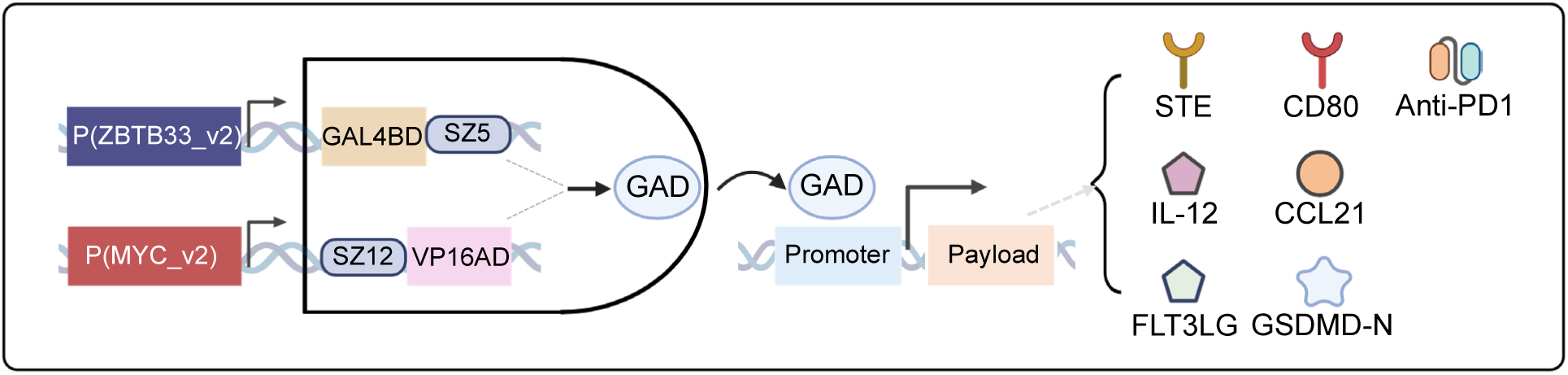
Schematic of the cancer-restricted AND-gate design. Two TREND-derived synthetic enhancers independently drive GAL4BD and VP16AD expression; coiled-coil pairing reconstitutes a synthetic transcription factor (GAD) that activates downstream therapeutic payloads, enabling cancer-selective expression of multiple immune effectors.

**Figure S9.**
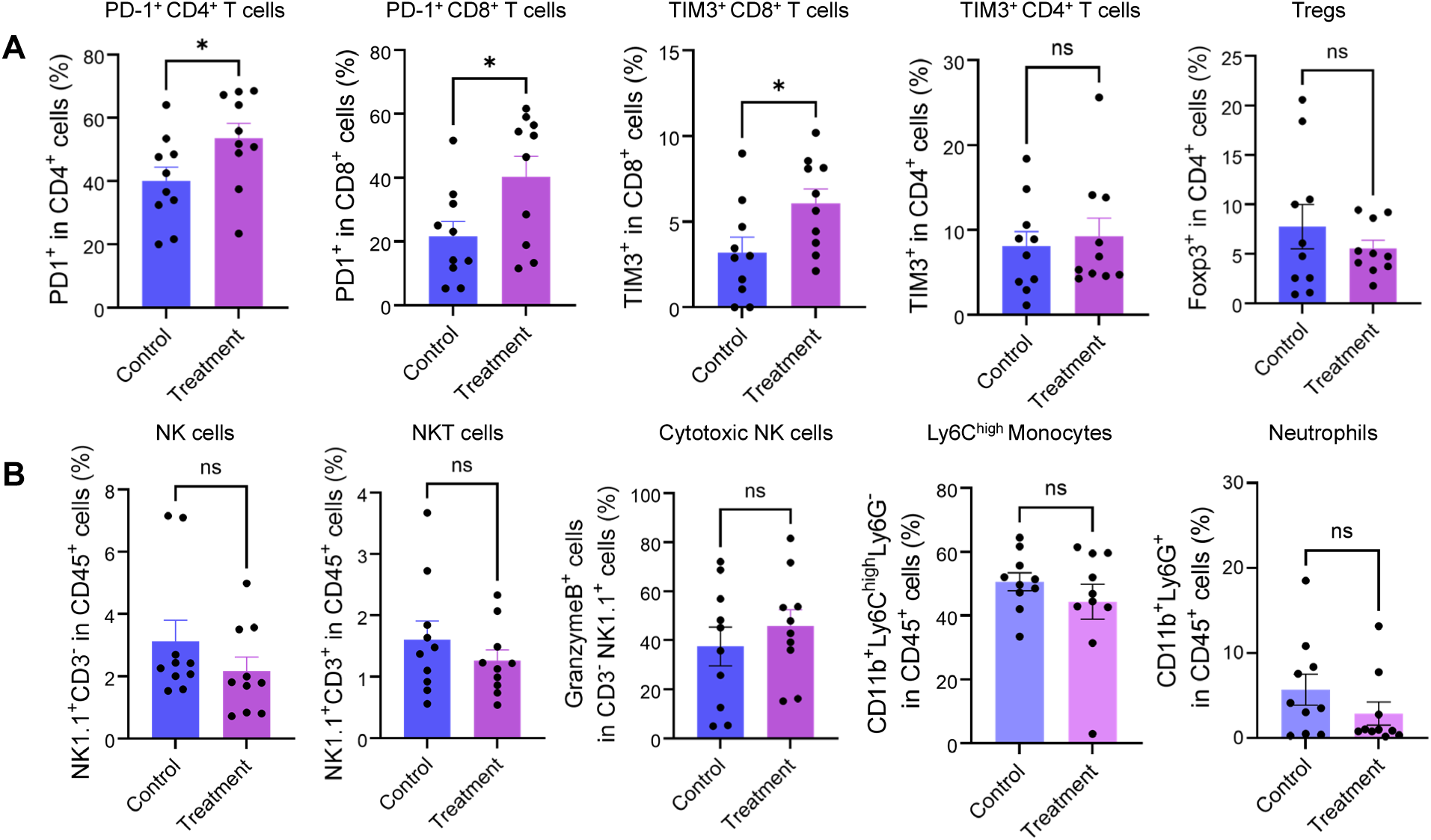
Characterization of tumor microenvironment (TME) immune composition following treatment with the AND-gate circuit expressing the S8IP output combination. (A) Flow-cytometric analysis of T-cell subsets in the peritoneal TME, showing proportion of PD-1^+^ CD4⁺ T cells, PD-1^+^ CD8^+^ T cells, TIM-3^+^ CD8⁺ T cells, TIM-3^+^ CD4^+^ T cells, and Tregs. (B) Proportion of innate immune populations including NK cells, NKT cells, cytotoxic NK cells, Ly6C^high^ monocytes and neutrophils. Data represent mean ± s.e.m. from 10 mice per group. Statistical significance was determined by unpaired two-tailed t-tests.

**Figure S10.**
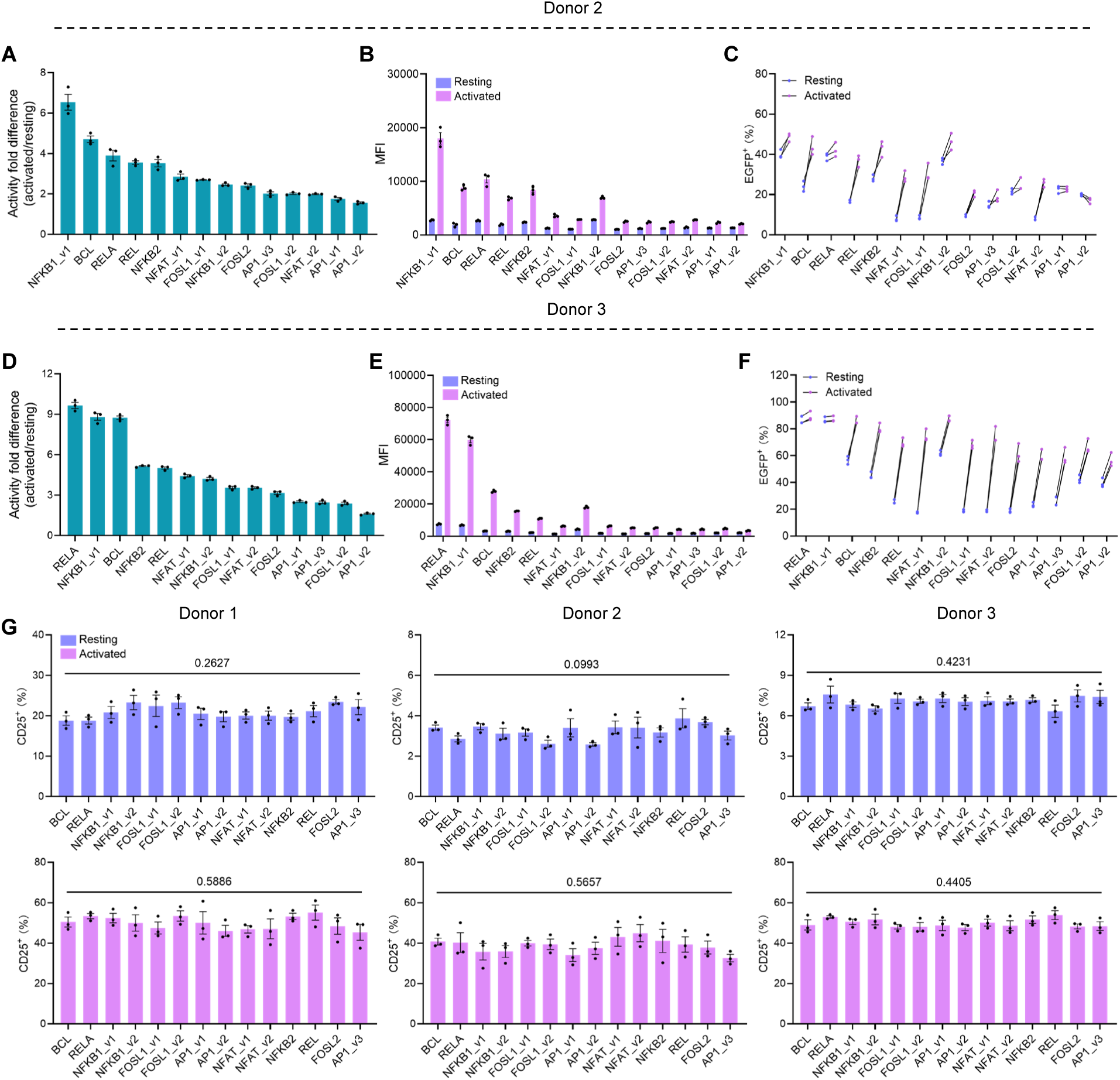
Validation of activation-responsive synthetic enhancers in primary human T cells. (A and D) Fold change in reporter activity (activated/resting) for all tested enhancers, identifying candidates responsive to T-cell activation. (B and E) Raw MFI of EGFP expression driven by individual enhancers under activated and resting conditions. (C and F) Percentage of EGFP⁺ T cells under activated versus resting conditions. (G) Proportion of CD25^+^ T cells transduced with individual enhancers under resting and activated conditions, summarized across three independent donors. Data represent mean ± s.e.m. from three independent experiments. Statistical significance was determined by unpaired two-tailed t-tests or one-way ANOVA.

**Figure S11.**
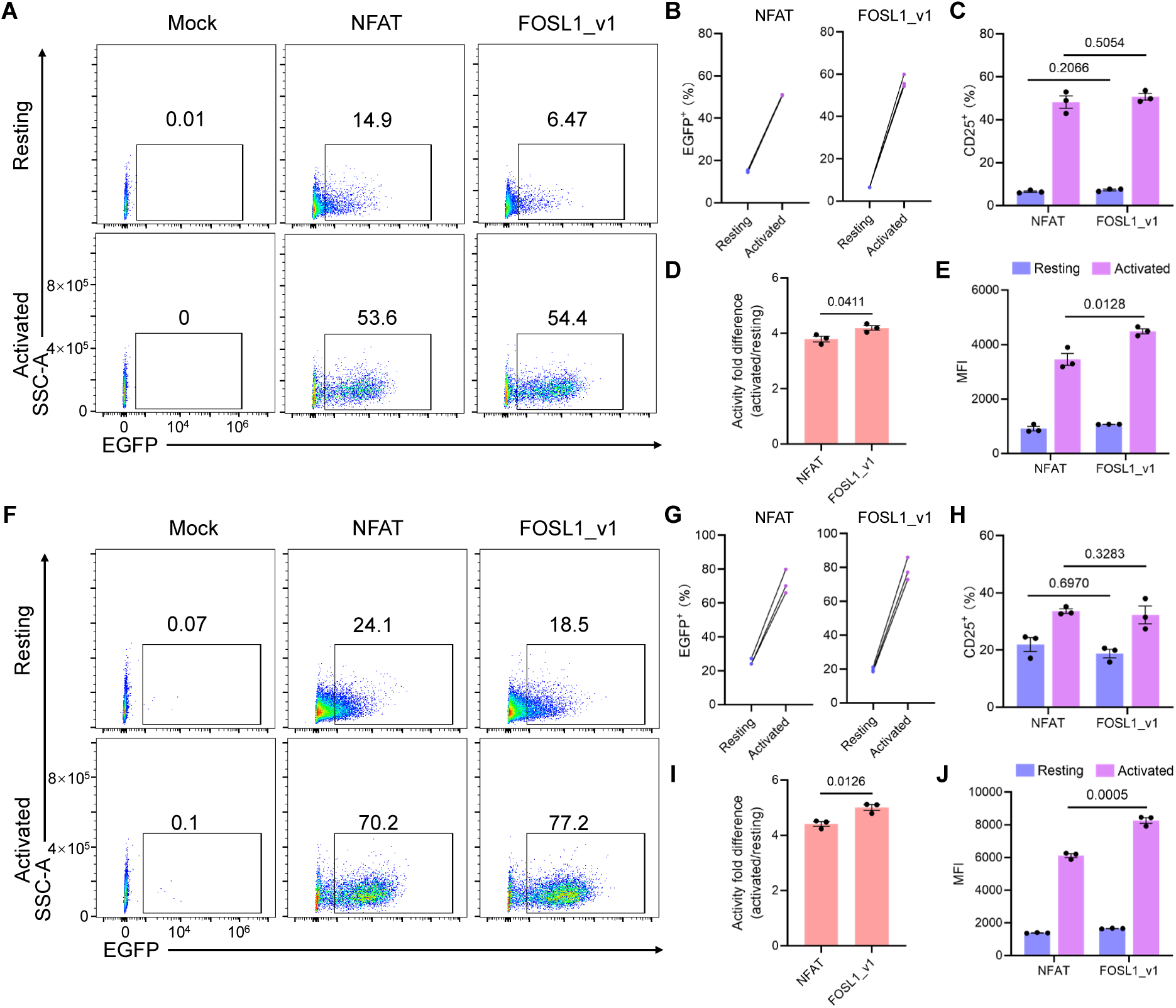
Comparison of NFAT and FOSL1-motif-based synthetic enhancer in T cells. (A and F) Representative flow plots showing EGFP expression driven by NFAT and FOSL1_v1 enhancers in resting and activated T cells. (B and G) Quantification of EGFP⁺ cell percentages from panels (A) and (F). (C and H) Proportion of CD25⁺ T cells transduced with NFAT or FOSL1_v1 enhancers under resting and activated conditions. (D and I) Fold change in reporter activity (activated relative to resting) for NFAT and FOSL1_v1 enhancers. (E and J) MFI of EGFP expression for NFAT and FOSL1_v1 enhancers in activated and resting T cells. Panels (A to E) and (F to J) show data from two different donors. Data are mean ± s.e.m. from three independent experiments. Statistical significance was determined by unpaired two-tailed t-tests.

## Notes

### Summary of Updates

This revised version includes new results and text revisions that strengthen the manuscript.

## Reference

1. Levo, M., and Segal, E. (2014). In pursuit of design principles of regulatory sequences. Nat Rev Genet 15, 453–468. 10.1038/nrg3684.

2. Jolma, A., Yan, J., Whitington, T., Toivonen, J., Nitta, K.R., Rastas, P., Morgunova, E., Enge, M., Taipale, M., Wei, G., et al. (2013). DNA-binding specificities of human transcription factors. Cell 152, 327–339. 10.1016/j.cell.2012.12.009.

3. Lambert, S.A., Jolma, A., Campitelli, L.F., Das, P.K., Yin, Y., Albu, M., Chen, X., Taipale, J., Hughes, T.R., and Weirauch, M.T. (2018). The Human Transcription Factors. Cell 172, 650–665. 10.1016/j.cell.2018.01.029.

4. Spitz, F., and Furlong, E.E. (2012). Transcription factors: from enhancer binding to developmental control. Nat Rev Genet 13, 613–626. 10.1038/nrg3207.

5. Butterfield, G.L., Reisman, S.J., Iglesias, N., and Gersbach, C.A. (2025). Gene regulation technologies for gene and cell therapy. Mol Ther 33, 2104–2122. 10.1016/j.ymthe.2025.04.004.

6. Chen, A.X.Y., Yap, K.M., Kim, J.S., Sek, K., Huang, Y.K., Dunbar, P.A., Wiebking, V., Armitage, J.D., Munoz, I., Todd, K.L., et al. (2025). Rewiring endogenous genes in CAR T cells for tumour-restricted payload delivery. Nature 644, 241–251. 10.1038/s41586-025-09212-7.

7. Nissim, L., Wu, M.R., Pery, E., Binder-Nissim, A., Suzuki, H.I., Stupp, D., Wehrspaun, C., Tabach, Y., Sharp, P.A., and Lu, T.K. (2017). Synthetic RNA-Based Immunomodulatory Gene Circuits for Cancer Immunotherapy. Cell 171, 1138–1150 e1115. 10.1016/j.cell.2017.09.049.

8. Gordon, M.G., Inoue, F., Martin, B., Schubach, M., Agarwal, V., Whalen, S., Feng, S., Zhao, J., Ashuach, T., Ziffra, R., et al. (2020). lentiMPRA and MPRAflow for high-throughput functional characterization of gene regulatory elements. Nat Protoc 15, 2387–2412. 10.1038/s41596-020-0333-5.

9. Liu, Y., Yu, S., Dhiman, V.K., Brunetti, T., Eckart, H., and White, K.P. (2017). Functional assessment of human enhancer activities using whole-genome STARR-sequencing. Genome Biol 18, 219. 10.1186/s13059-017-1345-5.

10. Agarwal, V., Inoue, F., Schubach, M., Penzar, D., Martin, B.K., Dash, P.M., Keukeleire, P., Zhang, Z., Sohota, A., Zhao, J., et al. (2025). Massively parallel characterization of transcriptional regulatory elements. Nature 639, 411–420. 10.1038/s41586-024-08430-9.

11. Fromel, R., Ruhle, J., Bernal Martinez, A., Szu-Tu, C., Pacheco Pastor, F., Martinez-Corral, R., and Velten, L. (2025). Design principles of cell-state-specific enhancers in hematopoiesis. Cell 188, 3202–3218 e3221. 10.1016/j.cell.2025.04.017.

12. Kosicki, M., Laboy Cintron, D., Keukeleire, P., Schubach, M., Page, N.F., Georgakopoulos-Soares, I., Akiyama, J.A., Plajzer-Frick, I., Novak, C.S., Kato, M., et al. (2025). Massively parallel reporter assays and mouse transgenic assays provide correlated and complementary information about neuronal enhancer activity. Nat Commun 16, 4786. 10.1038/s41467-025-60064-1.

13. Mauduit, D., Taskiran, II, Minnoye, L., de Waegeneer, M., Christiaens, V., Hulselmans, G., Demeulemeester, J., Wouters, J., and Aerts, S. (2021). Analysis of long and short enhancers in melanoma cell states. Elife 10. 10.7554/eLife.71735.

14. de Almeida, B.P., Schaub, C., Pagani, M., Secchia, S., Furlong, E.E.M., and Stark, A. (2024). Targeted design of synthetic enhancers for selected tissues in the Drosophila embryo. Nature 626, 207–211. 10.1038/s41586-023-06905-9.

15. Gosai, S.J., Castro, R.I., Fuentes, N., Butts, J.C., Mouri, K., Alasoadura, M., Kales, S., Nguyen, T.T.L., Noche, R.R., Rao, A.S., et al. (2024). Machine-guided design of cell-type-targeting cis-regulatory elements. Nature 634, 1211–1220. 10.1038/s41586-024-08070-z.

16. Taskiran, II, Spanier, K.I., Dickmanken, H., Kempynck, N., Pancikova, A., Eksi, E.C., Hulselmans, G., Ismail, J.N., Theunis, K., Vandepoel, R., et al. (2024). Cell-type-directed design of synthetic enhancers. Nature 626, 212–220. 10.1038/s41586-023-06936-2.

17. Wu, M.R., Nissim, L., Stupp, D., Pery, E., Binder-Nissim, A., Weisinger, K., Enghuus, C., Palacios, S.R., Humphrey, M., Zhang, Z., et al. (2019). A high-throughput screening and computation platform for identifying synthetic promoters with enhanced cell-state specificity (SPECS). Nat Commun 10, 2880. 10.1038/s41467-019-10912-8.

18. Shannon, P., and Richards, M. (2025). MotifDb: An Annotated Collection of Protein-DNA Binding Sequence Motifs (Bioconductor).

19. Hu, S., Xie, Z., Onishi, A., Yu, X., Jiang, L., Lin, J., Rho, H.S., Woodard, C., Wang, H., Jeong, J.S., et al. (2009). Profiling the human protein-DNA interactome reveals ERK2 as a transcriptional repressor of interferon signaling. Cell 139, 610–622. 10.1016/j.cell.2009.08.037.

20. Reddy, J., Fonseca, M.A.S., Corona, R.I., Nameki, R., Segato Dezem, F., Klein, I.A., Chang, H., Chaves-Moreira, D., Afeyan, L.K., Malta, T.M., et al. (2021). Predicting master transcription factors from pan-cancer expression data. Sci Adv 7, eabf6123. 10.1126/sciadv.abf6123.

21. D’Alessio, A.C., Fan, Z.P., Wert, K.J., Baranov, P., Cohen, M.A., Saini, J.S., Cohick, E., Charniga, C., Dadon, D., Hannett, N.M., et al. (2015). A Systematic Approach to Identify Candidate Transcription Factors that Control Cell Identity. Stem Cell Reports 5, 763–775. 10.1016/j.stemcr.2015.09.016.

22. Kribelbauer-Swietek, J.F., Pushkarev, O., Gardeux, V., Faltejskova, K., Russeil, J., van Mierlo, G., and Deplancke, B. (2024). Context transcription factors establish cooperative environments and mediate enhancer communication. Nat Genet 56, 2199–2212. 10.1038/s41588-024-01892-7.

23. Burgess, D.J. (2016). Regulatory elements: Putting enhancers into context. Nat Rev Genet 17, 377. 10.1038/nrg.2016.74.

24. Cheng, J.K., and Alper, H.S. (2016). Transcriptomics-Guided Design of Synthetic Promoters for a Mammalian System. ACS Synth Biol 5, 1455–1465. 10.1021/acssynbio.6b00075.

25. Shin, H.Y., Yang, W., Lee, E.J., Han, G.H., Cho, H., Chay, D.B., and Kim, J.H. (2018). Establishment of five immortalized human ovarian surface epithelial cell lines via SV40 T antigen or HPV E6/E7 expression. PLoS One 13, e0205297. 10.1371/journal.pone.0205297.

26. Sadowski, I., Ma, J., Triezenberg, S., and Ptashne, M. (1988). GAL4-VP16 is an unusually potent transcriptional activator. Nature 335, 563–564. 10.1038/335563a0.

27. Reinke, A.W., Grant, R.A., and Keating, A.E. (2010). A synthetic coiled-coil interactome provides heterospecific modules for molecular engineering. J Am Chem Soc 132, 6025–6031. 10.1021/ja907617a.

28. Das, A.T., Tenenbaum, L., and Berkhout, B. (2016). Tet-On Systems For Doxycycline-inducible Gene Expression. Curr Gene Ther 16, 156–167. 10.2174/1566523216666160524144041.

29. Iyer, S., Zhang, S., Yucel, S., Horn, H., Smith, S.G., Reinhardt, F., Hoefsmit, E., Assatova, B., Casado, J., Meinsohn, M.C., et al. (2021). Genetically Defined Syngeneic Mouse Models of Ovarian Cancer as Tools for the Discovery of Combination Immunotherapy. Cancer Discov 11, 384–407. 10.1158/2159-8290.CD-20-0818.

30. Anderson, K.G., Stromnes, I.M., and Greenberg, P.D. (2017). Obstacles Posed by the Tumor Microenvironment to T cell Activity: A Case for Synergistic Therapies. Cancer Cell 31, 311–325. 10.1016/j.ccell.2017.02.008.

31. Raskov, H., Orhan, A., Christensen, J.P., and Gogenur, I. (2021). Cytotoxic CD8(+) T cells in cancer and cancer immunotherapy. Br J Cancer 124, 359–367. 10.1038/s41416-020-01048-4.

32. Zhou, S.J., Liu, M.G., Ren, F., Meng, X.J., and Yu, J.M. (2021). The landscape of bispecific T cell engager in cancer treatment. BIOMARKER RESEARCH 9, 38. 10.1186/s40364-021-00294-9.

33. Liao, K.W., Chen, B.M., Liu, T.B., Tzou, S.C., Lin, Y.M., Lin, K.F., Su, C.I., and Roffler, S.R. (2003). Stable expression of chimeric anti-CD3 receptors on mammalian cells for stimulation of antitumor immunity. Cancer Gene Ther 10, 779–790. 10.1038/sj.cgt.7700637.

34. Powell, M.D., Read, K.A., Sreekumar, B.K., Jones, D.M., and Oestreich, K.J. (2019). IL-12 signaling drives the differentiation and function of a T(H)1-derived T(FH1)-like cell population. Sci Rep 9, 13991. 10.1038/s41598-019-50614-1.

35. Salmon, H., Idoyaga, J., Rahman, A., Leboeuf, M., Remark, R., Jordan, S., Casanova-Acebes, M., Khudoynazarova, M., Agudo, J., Tung, N., et al. (2016). Expansion and Activation of CD103+ Dendritic Cell Progenitors at the Tumor Site Enhances Tumor Responses to Therapeutic PD-L1 and BRAF Inhibition. IMMUNITY 44, 924–938. 10.1016/j.immuni.2016.03.012.

36. Lin, Y., Sharma, S., and John, M.S. (2014). CCL21 Cancer Immunotherapy. Cancers (Basel) 6, 1098–1110. 10.3390/cancers6021098.

37. Chen, R., Zou, J., Liu, J., Kang, R., and Tang, D. (2025). DAMPs in the immunogenicity of cell death. Mol Cell 85, 3874–3889. 10.1016/j.molcel.2025.09.007.

38. Patel, S.A., and Minn, A.J. (2018). Combination Cancer Therapy with Immune Checkpoint Blockade: Mechanisms and Strategies. IMMUNITY 48, 417–433. 10.1016/j.immuni.2018.03.007.

39. Han, J., Wang, S., Fang, W., Yang, Y., Zhou, R., Zhang, Y., Yu, J., Tang, R., Liu, Z., and Gu, Z. (2025). Long-acting IL-2 release from pressure-fused biomineral tablets promotes antitumor immune response. Nat Cancer 6, 1384–1399. 10.1038/s43018-025-00993-4.

40. Kramer, E.D., and Abrams, S.I. (2020). Granulocytic Myeloid-Derived Suppressor Cells as Negative Regulators of Anticancer Immunity. Front Immunol 11, 1963. 10.3389/fimmu.2020.01963.

41. Jeong, J., Suh, Y., and Jung, K. (2019). Context Drives Diversification of Monocytes and Neutrophils in Orchestrating the Tumor Microenvironment. Front Immunol 10, 1817. 10.3389/fimmu.2019.01817.

42. Zhang, L., Kerkar, S.P., Yu, Z., Zheng, Z., Yang, S., Restifo, N.P., Rosenberg, S.A., and Morgan, R.A. (2011). Improving adoptive T cell therapy by targeting and controlling IL-12 expression to the tumor environment. Mol Ther 19, 751–759. 10.1038/mt.2010.313.

43. Zhang, L., Morgan, R.A., Beane, J.D., Zheng, Z., Dudley, M.E., Kassim, S.H., Nahvi, A.V., Ngo, L.T., Sherry, R.M., Phan, G.Q., et al. (2015). Tumor-infiltrating lymphocytes genetically engineered with an inducible gene encoding interleukin-12 for the immunotherapy of metastatic melanoma. Clin Cancer Res 21, 2278–2288. 10.1158/1078-0432.CCR-14-2085.

44. Liu, Y., Di, S.M., Shi, B.Z., Zhang, H.H., Wang, Y., Wu, X.Q., Luo, H., Wang, H.M., Li, Z.H., and Jiang, H. (2019). Armored Inducible Expression of IL-12 Enhances Antitumor Activity of Glypican-3-Targeted Chimeric Antigen Receptor-Engineered T Cells in Hepatocellular Carcinoma. JOURNAL OF IMMUNOLOGY 203, 198–207. 10.4049/jimmunol.1800033.

45. Levine, B.L., Humeau, L.M., Boyer, J., MacGregor, R.R., Rebello, T., Lu, X., Binder, G.K., Slepushkin, V., Lemiale, F., Mascola, J.R., et al. (2006). Gene transfer in humans using a conditionally replicating lentiviral vector. Proc Natl Acad Sci U S A 103, 17372–17377. 10.1073/pnas.0608138103.

46. Scholler, J., Brady, T.L., Binder-Scholl, G., Hwang, W.T., Plesa, G., Hege, K.M., Vogel, A.N., Kalos, M., Riley, J.L., Deeks, S.G., et al. (2012). Decade-Long Safety and Function of Retroviral-Modified Chimeric Antigen Receptor T Cells. SCIENCE TRANSLATIONAL MEDICINE 4, 132ra53. 10.1126/scitranslmed.3003761.

47. Teixeira, A.P., and Fussenegger, M. (2024). Synthetic Gene Circuits for Regulation of Next-Generation Cell-Based Therapeutics. Adv Sci (Weinh) 11, e2309088. 10.1002/advs.202309088.

48. Bragdon, M.D.J., Patel, N., Chuang, J., Levien, E., Bashor, C.J., and Khalil, A.S. (2023). Cooperative assembly confers regulatory specificity and long-term genetic circuit stability. Cell 186, 3810–3825 e3818. 10.1016/j.cell.2023.07.012.

49. Liu, Y., Zeng, Y., Liu, L., Zhuang, C., Fu, X., Huang, W., and Cai, Z. (2014). Synthesizing AND gate genetic circuits based on CRISPR-Cas9 for identification of bladder cancer cells. Nat Commun 5, 5393. 10.1038/ncomms6393.

50. Guo, T., Ma, D., and Lu, T.K. (2022). Sense-and-Respond Payload Delivery Using a Novel Antigen-Inducible Promoter Improves Suboptimal CAR-T Activation. ACS Synth Biol 11, 1440–1453. 10.1021/acssynbio.1c00236.

51. Bonfá, G., Martino, G., Sellitto, A., Rinaldi, A., Tedeschi, F., Caliendo, F., Melchiorri, L., Perna, D., Starkey, F., Nikolados, E., et al. (2025). Design of novel synthetic promoters 1 to tune gene expression in T cells. bioaxiv. 10.1101/2025.01.09.632034.

52. Inoue, F., and Ahituv, N. (2015). Decoding enhancers using massively parallel reporter assays. Genomics 106, 159–164. 10.1016/j.ygeno.2015.06.005.

53. Lim, F., Solvason, J.J., Ryan, G.E., Le, S.H., Jindal, G.A., Steffen, P., Jandu, S.K., and Farley, E.K. (2024). Affinity-optimizing enhancer variants disrupt development. Nature 626, 151–159. 10.1038/s41586-023-06922-8.

54. Kheradpour, P., and Kellis, M. (2014). Systematic discovery and characterization of regulatory motifs in ENCODE TF binding experiments. Nucleic Acids Res 42, 2976–2987. 10.1093/nar/gkt1249.

